# The Sulfated PSY Peptide Negatively Regulates Receptor Kinase Activity to Promote Growth

**DOI:** 10.64898/2026.07.25.740709

**Authors:** Devin V. Tulio, Alexandra M. Shigenaga, Dawn Lim, Artur Teixeira de Araujo, Shu-Zon Wu, Pamela C. Ronald, Magdalena Bezanilla

**Affiliations:** Department of Biological Sciences, Dartmouth College, Hanover, NH 03755; Department of Plant Pathology and the Genome Center, University of California, Davis, CA 95616

**Author notes:** These authors contributed equally (Devin V. Tulio, Alexandra M. Shigenaga).

## Abstract

Complex signaling pathways organize cell expansion and proliferation across cells to pattern tissues and organs in plants. The sulfotyrosine peptide hormone family, <u>P</u>LANT PEPTIDE CONTAINING <u>S</u>ULFATED T<u>Y</u>ROSINE (PSY), contributes to these processes. We identified two plasma membrane-localized receptors, PSYR1 and PSYR2, that are necessary for PSY signaling and regulate growth in *Physcomitrium patens.* Membrane-associated PSYRs accumulate to high levels in a mutant lacking TYROSYL PROTEIN SULFOTRANSFERASE (TPST). Given that a *tpst* null mutant (*Δtpst*) is impaired in sulfation, this suggests that in the absence of sulfated peptides, PSYRs accumulate on the membrane. A null mutant of the PSY receptors, *Δpsyr1/*2, showed increased growth and was epistatic to *Δtpst,* suppressing defects in gametophore formation and early senescence. The transcriptional profiles comparing wild type to *Δpsyr1/2* and *Δpsyr1/2/Δtpst* showed 25 to 30 differentially expressed genes between the receptor null mutants and wild type, with a common signature of cell wall remodeling and stress responses. Similarly, a PSYR1 kinase-inactive mutation rescued *Δtpst* and relieved the accumulation of membrane-associated PSYRs. In contrast, overexpression of PSYRs inhibited plant growth, with phenotypic severity correlating with the amount of overexpression. These data are consistent with a constitutive activation model in which membrane-associated PSYRs unbound to PSY serve to inhibit growth through an active kinase. In the presence of the PSY peptide, the kinase is inactivated, promoting growth and driving *PSY* expression. The relationship between growth-repressive PSYR kinase activity and growth-promoting PSYR kinase inactivation in *P. patens* serves as a model for optimizing plant growth and development.

**Significance Statement:** As sessile organisms, plant growth and development rely on the intricate integration of external and internal signals. Plants produce peptide hormones and those signals are read by leucine rich receptor like kinases (LRR-RLKs), which normally are activated by binding to peptide. Taking advantage of the ability to generate point mutations directly in the genome, we show that PSY peptide binding leads to kinase inactivation, shifting the paradigm for peptide signaling in plants.

## Introduction

As sessile organisms, plant growth and development rely on the intricate integration of external and internal signals. Reading and deciphering these signals depends in part on plant receptor-like kinases (RLKs). The majority of RLKs across plant species belong to a subfamily that share an extracellular leucine-rich repeat (LRR) domain, known as LRR-RLKs. LRR-RLKs are highly conserved and mediate both developmental and stress responses^1–3^. Within the 19 subfamilies of LRR-RLKs, subfamilies X and XI are implicated in endogenous-peptide-mediated signaling^3,4^. A subset of receptors in subfamily X and XI recognize tyrosine-sulfated peptide hormones and influence growth, development, and immune response^3,5,6^. Four sulfated tyrosine peptide hormone families have been identified and characterized in higher plants including the <u>P</u>LANT PEPTIDE-CONTAINING <u>S</u>ULFATED T<u>Y</u>ROSINE (PSY) family, <u>P</u>HYTO<u>S</u>ULFO<u>K</u>INES (PSKs), <u>R</u>OOT MERISTEM <u>G</u>ROWTH <u>F</u>ACTORS (RGFs), and <u>C</u>ASPARIAN STRIP INTEGRITY <u>F</u>ACTORS (CIFs); all of which play roles in multiple aspects of plant development, ranging from seed and pollen development to Casparian strip formation, lateral root initiation, root apical meristem development, cell proliferation and division^7–18^.

The cognate receptors for each of these tyrosine-sulfated peptides have been identified in angiosperms: PSY receptors (PSYRs, also known as ROOT ELONGATION RECEPTOR KINASES/REKs), PSK receptors (PSKRs), RGF receptors (RGFRs, also known as RGF1-INSENSITIVE/RGIs), and the CIF receptors (GSO/SGN; GASSHO/SCHENGEN)^9,11,12,19–24^. However, identification of these receptors in other plant lineages is limited. Of particular interest are the PSY receptors^23,24^, which, in Arabidopsis, have a unique mode of action compared with the receptors for the other three sulfated peptide classes. In the absence of the PSY peptides, the AtPSYRs suppress growth-related pathways and activate transcription factor genes related to stress responses^23^. This PSYR-mediated ligand-deprivation-dependent activation system has not yet been investigated in other plant species.

In contrast to angiosperms, the spreading earth moss, *Physcomitrium patens,* genome encodes a single *PSY homolog and PSK homolog*^25^*, as well as two immune-responsive PSY-like peptides, PSYL1 and PSYL2*^26^. However, the PpPSK peptide previously identified^25^ lacks the conserved residue required for tyrosine sulfation, and is therefore unlikely to be sulfated^27^. To date, PSY peptides are the only tyrosine-sulfated peptide family predicted to be functional in *P. patens*^26,27^. Together, this highlights the usefulness of *P. patens* as an advantageous model plant system for studying tyrosine-sulfated peptide signaling.

Furthermore, although *P. patens* has a different body form from angiosperms, *P. patens* utilizes developmental principles universal to all plants, such as the establishment of proliferative stem cells and cell expansion-making *P. patens* an excellent model system for studying conserved mechanisms of plant development. The initial tissue establishing the *P. patens* plant emerges from a spore, protoplast or damaged tissue, comprising an essentially two-dimensional (2D) filamentous network. This tissue branches and expands at the tips of the filaments via tip growth and cell division of the apical stem cell. As protonemata ages, some branching events transition to three-dimensional (3D) growth, forming a bud that establishes a tetrahedral stem cell. Subsequently, diffuse cell expansion leads to formation of a shoot structure that is known as the gametophore.

One challenge for in-depth characterization of PSYR receptors arises from the presence of multiple paralogs of each sulfated tyrosine peptide family in the genomes of all angiosperms characterized to date. However, because there is only one enzyme that catalyzes the post-translational modification of sulfated tyrosine peptides, it has been possible to study how loss of sulfated peptides impacts growth and development across plant lineages. This enzyme, TYROSYL PROTEIN SULFOTRANSFERASE (TPST) is encoded by a single gene in Arabidopsis and *P. patens*. In Arabidopsis, TPST mediates sulfation of PSYs, PSKs, RGFs, and CIFs^6,28^. Thus, a *tpst* null mutant represents deficiency in sulfotyrosyl proteins including all the tyrosine-sulfated peptides. In Arabidopsis, *TPST* is highly expressed in the root apical meristem^29^ and a *tpst* null mutant has growth and developmental deficits, including dwarfism, stunted roots, reduced formation of flowers and higher order veins, as well as early senescence^29^. Strikingly, exogenous application of PSY1 to *tpst* partially restores root growth in Arabidopsis^10,30,31^, which is consistent with data demonstrating that overexpression of PSYs or exogenous application to wild-type plants causes root elongation in Arabidopsis and rice^9,14,30,31^. These studies have pointed towards PSY signaling as a critical regulator of root elongation in angiosperms. Similarily, null mutants in the sole copy of *P. patens TPST* (*Δtpst*) exhibit developmental deficits similar to those found in Arabidopsis, ranging from reduced plant size, defective gametophore development and expansion, and early senescence^27^. Addition of exogenous PSY peptide derived from either *P. patens* or Arabidopsis largely rescues these developmental deficits. Furthermore, similar to the angiosperm PSYs, the addition of *P. patens* PSY to both wild-type and *tpst-1* plants in Arabidopsis and wild-type rice seedlings promoted root elongation, providing strong evidence that PSY signaling is evolutionarily conserved^27^.

To date, PpTPST has been demonstrated to sulfate one peptide family, PSY^27^. Furthermore, given that *P. patens* encodes fewer LRR-RLKs compared to Arabidopsis and rice^3^, is easy to genome edit^32^, and has a predominantly haploid life cycle^32^, *P. patens* provides a simpler system to probe how tyrosine-sulfated peptide signaling controls growth and development. Here we report the identification and characterization of two putative *P. patens* PSY receptors (PpPSYR1 and PpPSYR2). Using a combination of genetics and live cell imaging, we found that the *P. patens* PSY receptors accumulate on the plasma membrane in plants that have lost the ability to produce tyrosine-sulfated peptides. This accumulation inhibits growth via an active kinase, and the severity of growth inhibition is related to the levels of kinase-active receptors on the membrane. Our results demonstrate that *P. patens* PSYRs employ a ligand-deprivation-dependent activation system as described for PSY signaling in Arabidopsis. In contrast to Arabidopsis, we provide further insight into the underlying mechanism of this unique peptide signaling pathway. We show that PSY-signaling fine-tunes plant growth and development in *P. patens* through negative feedback between membrane-associated PSYR kinase activity, which suppresses PSY-mediated growth processes, and transcriptional control of the *PSY* ligand, which inactivates PSYR kinase activity.

## Results

### Identification and characterization of PSY receptors in P. patens

Using the Arabidopsis PSY receptors AtPSYR1,2,3^23,24^ as references, we identified putative PSYR orthologs in *P. patens* from published LRR-RLK sub-family XI phylogenies^3,25^ in which two *P. patens* genes (Pp3c24_15950/XP_024364615 and Pp3c8_17530/XP_024381748) cluster with the corresponding AtPSYR clade. We confirmed orthology by a reciprocal best-hit BlastP test. The full-length protein sequences from AtPSYR1 (At1g17230), AtPSYR2 (At2g33170), and AtPSYR3 (At5g63930) were used as a BlastP query against the *P. patens* set (taxid:3218). All three AtPSYRs returned the same two *P. patens* proteins as their top hits genome-wide, with similar identity: Pp3c24_15950 (48.31/48.89/48.18% identity) and Pp3c8_17530 (48.09/47.34/47.79% identity). For the reciprocal search, we compared the full-length protein sequences encoded by Pp3c24_15950 and Pp3c8_17530 to the *A. thaliana* set (taxid: 3702). For both *P. patens* proteins, the best Arabidopsis hits were the AtPSYRs, which also yielded similar identity scores (Pp3c24_15950: 48.73/48.61/48.48%; Pp3c8_17530: 48.39/48.42/47.83% to AtPSYR1/2/3, respectively). Together, these results show a consistent pattern: all three AtPSYRs returned the same two *P. patens* proteins at comparable identity, rather than each matching a distinct moss protein. This indicates that the PpPSYR genes are co-orthologs of the AtPSYR clade rather than of any single paralog. This is expected if AtPSYRs arose from Arabidopsis-lineage duplications after divergence from *P. patens*, and further supports the basal placement of the PpPSYRs relative to the AtPSYR clade in the LRR-RLK XI phylogenies^3,25^. We therefore designated Pp3c24_15950 as PpPSYR1 and Pp3c8_17530 as PpPSYR2 (referred to as PSYR1 and PSYR2 hereafter in the text). Further support for these *P. patens* PSY receptor candidates comes from comparison to putative rice (*Oryza sativa*) PSY receptors (OsPSYR1/LOC_Os07g05740 and OsPSYR2/LOC_Os04g42700; numbering determined by reciprocal best hit to corresponding AtPSYR; Fig. S1A). The Arabidopsis, *P. patens*, and putative rice PSY receptors all contain the same domain structure (Fig. S1A).

PSYR1 is 1156 amino acids and PSYR2 is 1154 amino acids. Sequence alignment of the PSYR1 and PSYR2 ectodomains, based on the plant-specific LRR-domain^33^, revealed that similar to AtPSYR1, both PSYR1 and PSYR2 encode 26 LRRs (Fig. 1A). Conserved residues in the LRR-domains between the Arabidopsis, *P. patens,* and putative *O. sativa* PSYRs are highlighted in Fig. 1A and Fig. S1A. Based on previous studies with other sulfated-peptide receptors, specifically the Arabidopsis RGI/RGFR, the ectodomain “RxGG” motif is required to recognize the sulfated tyrosine of the Arabidopsis RGF1 peptide^22^. Based on the corresponding LRR position (LRR06) of the “RxGG” motif, the AtPSYRs and PpPSYRs share the “RxG” core but diverge at the fourth position (Fig. S1B). Because the “RxGG” motif is specific to the RGF1-RGFR interaction, this divergence is expected for receptors in the PSYR clade. However, there are a broader set of conserved RGFR residues that recognize the sulfated tyrosine such as the RGFR Asp-174, Arg-195, Gly-197, Gly-220, and Ala-222^22^, which are conserved across the Arabidopsis, *P. patens*, and putative rice PSYRs (Fig. S1B) and all of which were shown to bind the sulfated tyrosine residue in RGF^22^. The conservation of sulfotyrosine-binding residues, despite divergence from the RGFR specific “RxGG” motif, indicates that the *P. patens* PSYR1 and PSYR2 preserve structural features associated with sulfated-peptide binding and recognition.

**Figure 1.**
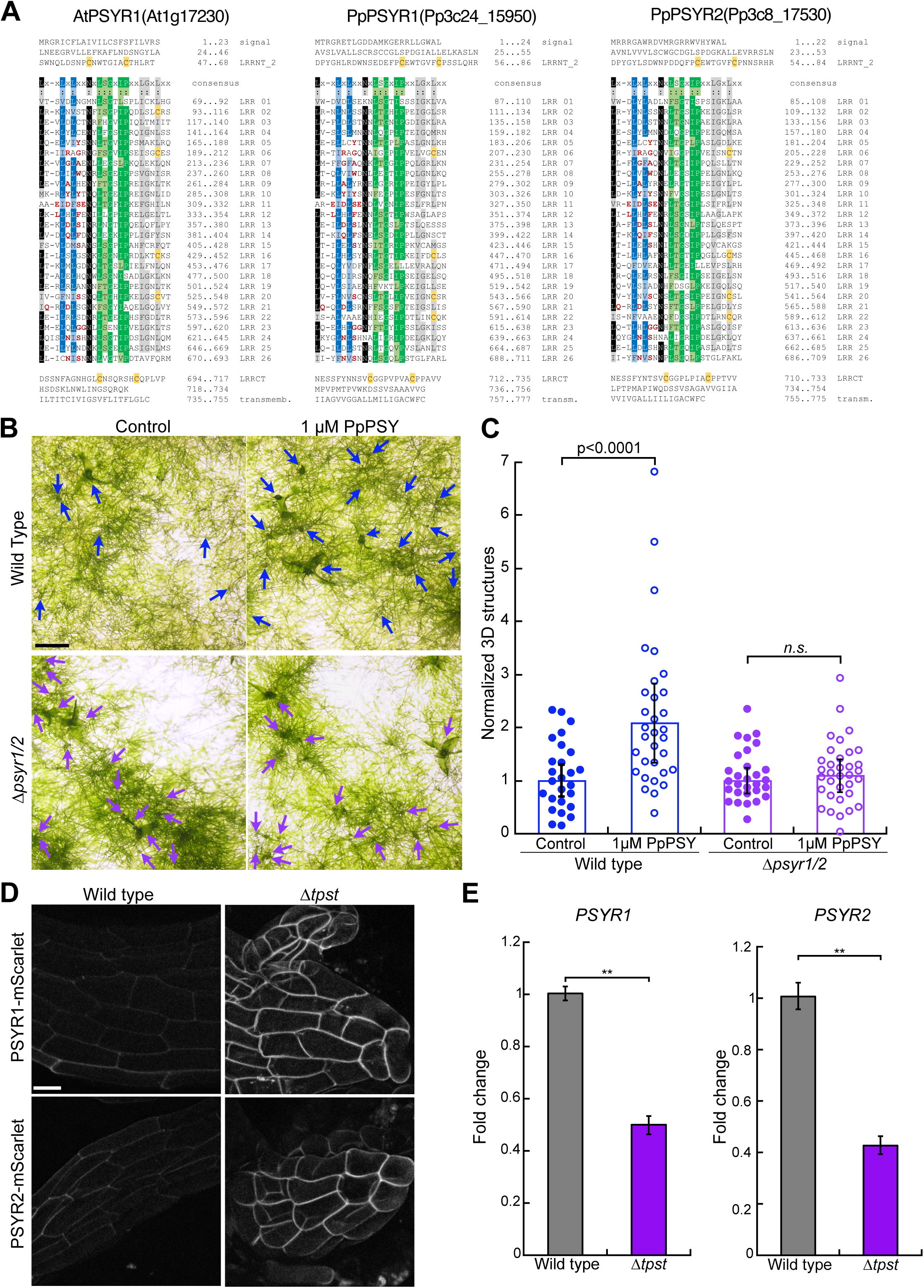
Identification and Characterization of *P. patens* PSY Receptors. A) Sequence alignments of the ectodomain (Signal peptide, LRR-cap, LRR-domain to transmembrane domain) for AtPSYR1, PpPSYR1, and PpPSYR2. Conserved residues across these three receptors and AtPSYR2 and AtPSYR3 are highlighted in red. The LRR-domains are organized based on the plant-specific LRR consensus (LxxLxLxxNxLsGxIPxxLGxLxx). B) *PSY* receptor null mutants are insensitive to PSY. Representative brightfield images of 12-day-old ground tissue on media supplemented with 1µM PpPSY. Arrows mark buds and gametophores in each image. Scale bar, 0.5 mm. Error bars represent standard error (n=3 separate trials). C) Quantification of the number of 3D structures (buds and gametophores) in control and peptide-treated tissue per field of view. The number of 3D structures is normalized to the average number of 3D structures in the control group for each line. Significance was calculated using a Kruskal-Wallis test. n.s., not significant. D) PSYRs localize to the plasma membrane and are enhanced in *Δtpst.* Maximum intensity projections of confocal z-stacks of PSYR1-mScarlet or PSYR2-mScarlet in an expanded gametophore phyllid (wild type) or aborted gametophore (*Δtpst*) dissected from approximately 4-week-old ground tissue. Scale bar, 20 µm. E) *PSYR1* and *PSYR2* transcript levels in approximately 4-week-old wild-type and *Δtpst* ground tissue. Significance was calculated using Welch’s t-test with a significance level of α=0.05. n.s., not significant.

If PSYR1 and PSYR2 are the receptors that bind to PSY, we would expect that a null mutant lacking both receptors should not respond to exogenous application of PSY. To test this, we used CRISPR-Cas9 mediated genome editing to generate a *psyr* null mutant (*Δpsyr1/2*) (Fig. S2, Table S1). We grew wild-type and *Δpsyr1/2* tissue on media supplemented with PpPSY. Since 1µM of PpPSY was sufficient to rescue *Δtpst*^27^, we tested this concentration on wild-type and *Δpsyr1/2* plants. After 12 days on peptide media, we found that wild type exhibited an increase in the number of buds and gametophores (Fig. 1B), forming on average 2.08 times more buds and gametophores compared to control conditions (Fig. 1C). In contrast, *Δpsyr1/*2 treated with peptide had similar numbers of buds and gametophores as the control condition (Fig. 1C). These data suggest that *Δpsyr1/2* is insensitive to PSY1, providing evidence that PSYR1 and PSYR2 respond to PSY1. Under control conditions, *Δpsyr1/*2 appeared to form more buds and gametophores than wild type (Fig. 1B), suggesting that *Δpsyr1/2* exhibits a constitutive PSY response.

To determine how peptides affect PSY receptor localization, we used CRISPR-mediated homology-directed repair (HDR) to endogenously tag the two *P. patens* PSY receptors (*PSYR1* and *PSYR2*). We fused sequences encoding mScarlet at the 3’ end of each gene just before the stop codon, which would ultimately position mScarlet in the cytoplasm (Fig. S3, Table S1). Under our imaging conditions, while we did not detect mScarlet fluorescence in protonemata, we did observe fluorescence in gametophores. Both PSYR1 and PSYR2 localized to the plasma membrane of expanding phyllids (Fig. 1D). However, in plants lacking sulfated peptides (*Δtpst*), the fluorescent signal was noticeably higher for both PSYR1 and PSYR2 compared to wild type (Fig. 1D). With PSYR localization outlining each cell, we found that wild-type cells were more uniform and generally were elongated polygons, whereas *Δtpst* cells were rounder and less oblate (Fig. 1D). Because the receptor genes were endogenously tagged, the fluorescence levels represent physiologically relevant protein levels. Interestingly, the increase in PSYR1 and PSYR2 fluorescent signals in *Δtpst* was not a result of higher transcript levels, with *Δtpst* having only 0.42 and 0.50 the amount of PSYR1 and PSYR2 transcript levels, respectively, compared to wild type (Fig. 1E). Therefore, these data suggest that the absence of sulfated peptides leads to receptor protein accumulation on the membrane independent of transcription.

### Loss of TPST in the psyr null mutant rescues gametophore development and expansion in Δtpst

If the defect in gametophore expansion in *Δtpst* results from increased levels of PSYR1 and PSYR2 on the membrane, we predicted that loss of the receptors from the membrane should rescue gametophore expansion. To test this, we generated a *tpst* null mutant in *Δpsyr1/2* (Fig. S2, Table S1). With only two PSY receptor genes in *P. patens*, *Δpsyr1/2* represents the complete loss of PSY receptor signaling. As predicted, both Δ*psyr1/2* and *Δpsyr1/2/Δtpst* plants formed expanded gametophores, whereas *Δtpst* plants were dense and lacked expanded gametophores (Fig. 2A^27^). Interestingly, both *Δpsyr1/2* and *Δpsyr1/2/Δtpst* plants were less dense at the center of the plant and projected extensive protonemal tissue outwards past the main body of the plant (Fig. 2A). This increased two-dimensional (2D) expansion compared to wild type, resulted in an average area where *Δpsyr1/2* and *Δpsyr1/2/Δtpst* were 18% and 16% larger than wild type respectively, whereas *Δtpst* was 69.5% smaller than wild type (Fig. 2B). In addition to the formation of expanded gametophores, *Δpsyr1/2/Δtpst* plants did not senesce early, which is associated with loss of TPST function (Fig. 2A^27^).

**Figure 2.**
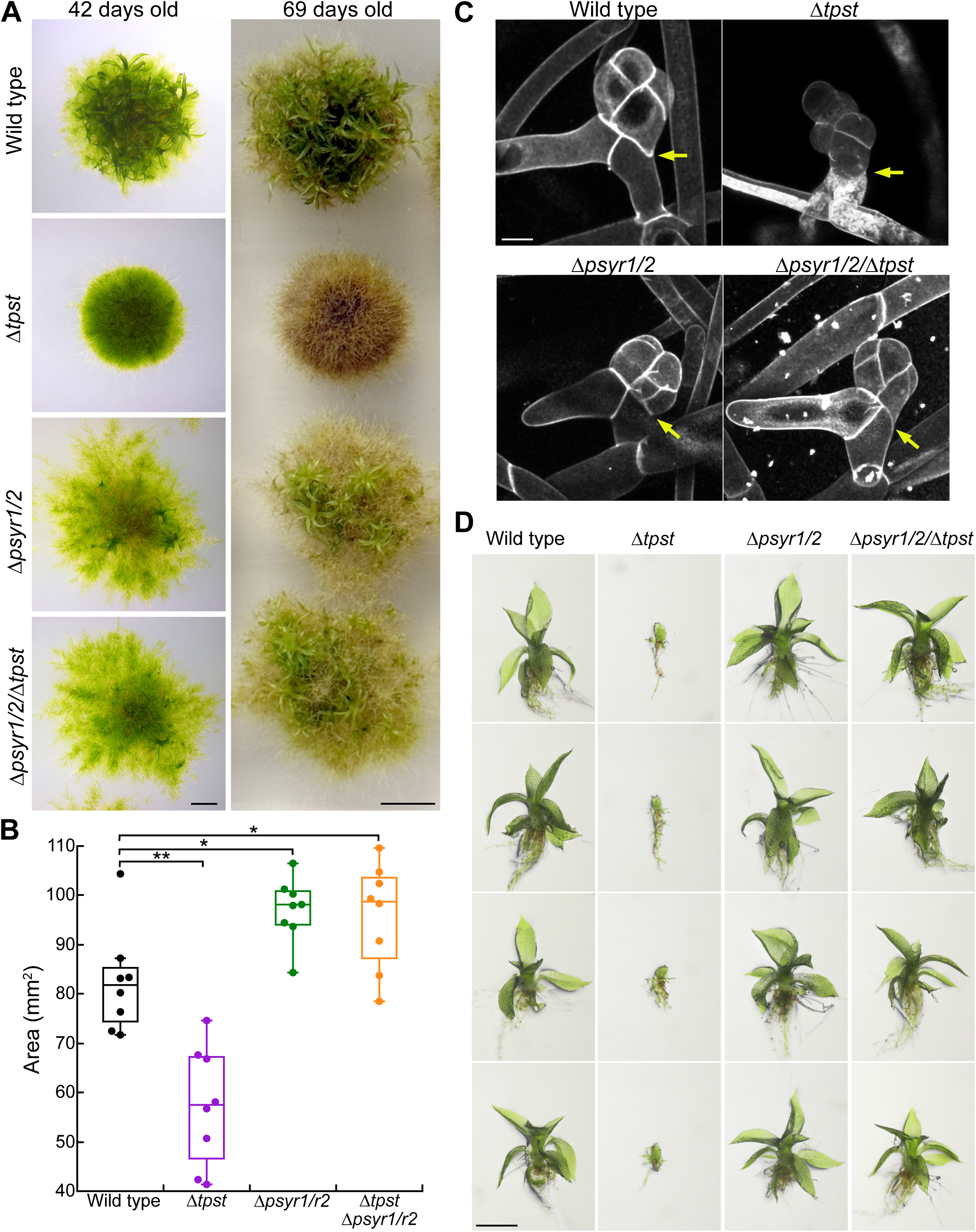
Loss of PSYR function rescues *Δtpst*. A) Representative brightfield images of 42-day-old wild-type, *Δtpst, Δpsyr1/2,* and *Δpsyr1/2/Δtpst* plants regenerated from protoplasts. Scale bar, 0.2 cm. Images on the right depict the same plants at 69-days-old. Scale bar, 0.5 cm. B) Quantification of plant area of 42-day-old plants. Significant differences determined by a Kruskal-Wallis test (α=0.05). **Wild type vs *Δtpst*, p= 0.0016; *Wild type vs *Δpsyr1/2*, p=0.0117; *Wild type vs *Δpsyr1/2/Δtpst*, p=0.0274. C) Representative maximum intensity projections of confocal z-stacks from propidium-iodide-stained buds. Yellow arrow denotes the first division in the bud initial. Scale bar, 20 µm. D) Representative extended depth of focus brightfield images of single gametophores dissected from 4-week-old ground tissue. Scale bar, 0.5 mm.

The *tpst* null mutant is defective in bud development, typically marked early on by a non-oblique first division as well as a reduction in cell expansion^27^. Both *Δpsyr1/2* and *Δpsyr1/2/Δtpst* did not show any abnormalities in early bud development with a normal oblique first division (Fig. 2C). Moreover, individual *Δpsyr1/2* and *Δpsyr1/2/Δtpst* expanded gametophores were unremarkable compared to wild type (Fig. 2D). Taken together, these data demonstrate that loss of PSYRs rescues gametophore development and expansion and rescues early senescence observed in *Δtpst*.

### Transcriptional landscape of tpst and psy receptor null mutants

To ask how sulfated-peptide signaling shapes gene expression in *P. patens*, we compared the transcriptomes of wild type, *Δtpst*, *Δpsyr1/2,* and *Δpsyr1/2/Δtpst*. The *Δpsyr1/2* mutant represents the complete loss of PSY receptor signaling, while the *Δpsyr1/2/Δtpst* mutant represents the simultaneous loss of receptor signaling and tyrosine sulfation, and is therefore the epistatic double perturbation. Three biological replicates were analyzed per genotype.

Differentially expressed genes (DEGs) were distributed unevenly across comparisons (Fig. 3A, Table S2). Loss of the PSY receptors had little effect on the transcriptome. Only 25 genes were differentially expressed between wild type and *Δpsyr1/2* and 30 between wild-type and *Δpsyr1/2/Δtpst*. In contrast, loss of TPST altered 271 genes relative to wild type, and the largest differences of all were observed when comparing the *Δtpst* transcriptional landscape with the receptor-null lines (∼1,800 genes). The larger DEG count seen in the *Δtpst*-referenced contrasts is due to the reference itself: the *tpst* null mutant differs from wild type at 271 genes, so any comparison that uses *Δtpst* as a reference inherits a large baseline DEG count. However, the identity and direction of individual genes in these comparisons remain informative. Wild type, *Δpsyr1/2* and the *Δpsyr1/2/Δtpst* were mutually similar, whereas Δ*tpst* was the clear transcriptional outlier (Fig. S4A, Table S2). At the transcriptome level, preventing ligand sulfation by knocking out *TPST* altered far more genes than removing the *PSY* receptors (wild type vs *Δtpst*: 271 DEGs; wild type vs *Δpsyr1/2*: 25 DEGs). To validate the RNA-Seq data, we measured the expression of four candidate genes by quantitative RT-PCR (RT-qPCR) (Pp3c3_36680, Pp3c2_1870, Pp3c8_7340, and Pp3c9_1620; highlighted in Fig. S4A). In wild type, *Δtpst, Δpsyr1/2,* and *Δpsyr1/2/Δtpst*, the RT-qPCR data confirmed the differential expression detected by RNA-Seq analysis (Fig. S4B).

**Figure 3:**
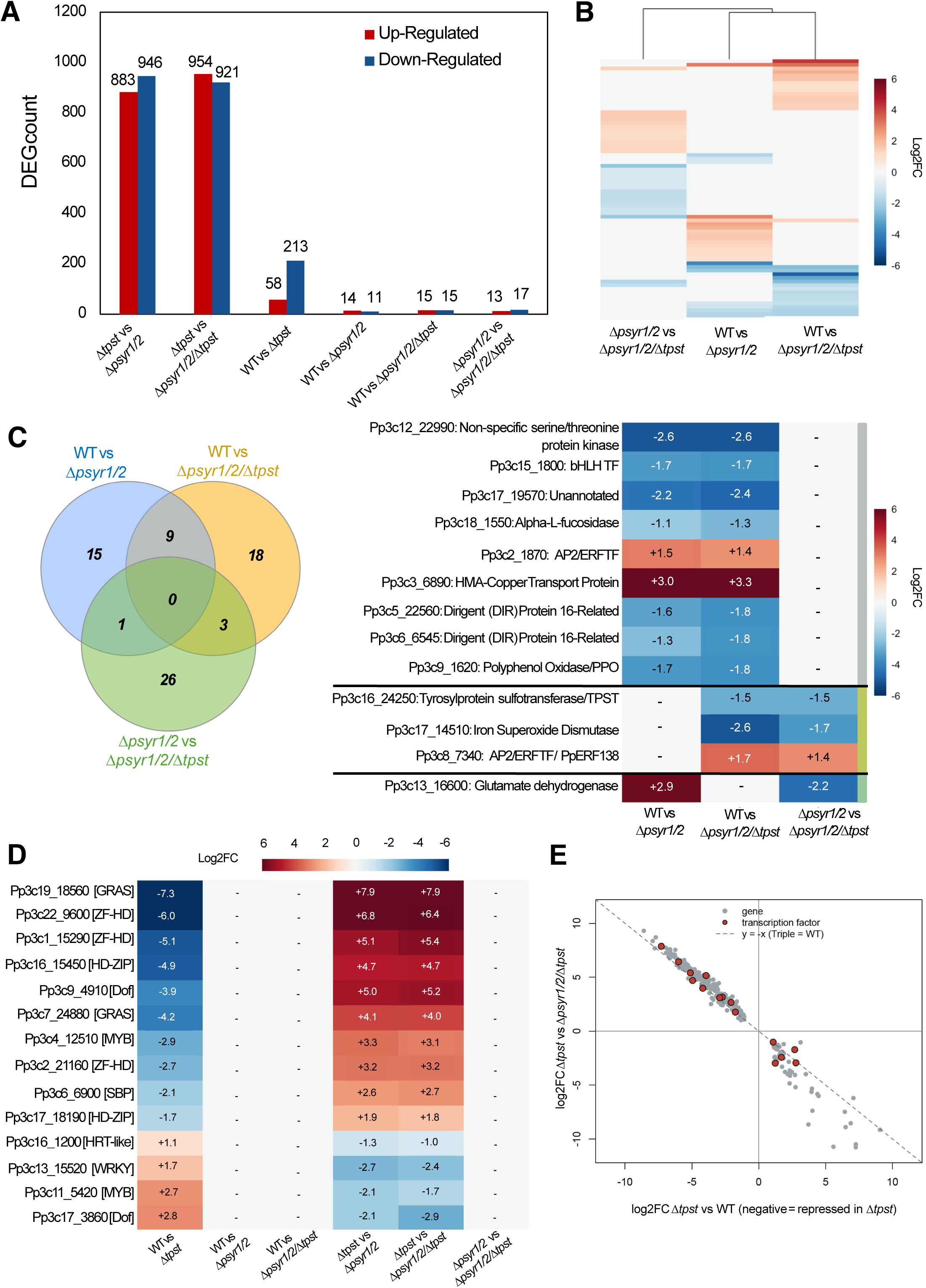
Genome-wide transcriptional responses across the sulfation-null and receptor-null comparisons. A) Number of differentially expressed genes in each pairwise comparison, separated into up- (red) and down-regulated (blue) genes; counts are shown above and below each bar (direction relative to the second-named genotype). B) Heatmap of log₂ fold-changes for the genes differentially expressed in at least one of the three comparisons (columns: Wild type (WT) vs *Δpsyr1/2*; WT vs *Δpsyr1/2/Δtpst*; *Δpsyr1/2* vs *Δpsyr1/2/Δtpst*); rows clustered; red = higher in the second-named genotype; blue = lower. C) Venn Diagram (left) shows number of differentially expressed genes overlapping among the three comparisons. Nine genes are shared by the two receptor-null lines, and the genes distinguishing *Δpsyr1/2/Δtpst* from *Δpsyr1/2* define a TPST-dependent, PSY receptor-independent set. Log₂ fold-changes (right) of the nine shared *Δpsyr1/2* genes and the *TPST*-dependent/*PSYR*-independent genes across the three comparisons; color as in (B); a dash (-) marks genes not differentially expressed in that comparison. Colors next to each set of genes correlates to the colors overlapping in the Venn Diagram. D) Log₂ fold-changes, across all six comparisons of the transcription factors within the PSY receptor-gated set; color and dashes as in (B and C). E) Loss of TPST produces a PSY receptor-gated, growth-repressive program. For genes differentially expressed in both comparisons (n = 262), fold-changes in WT vs *Δtpst* and *Δtpst/Δpsyr1/2* vs Δ*tpst* are anti-correlated (r = −0.984), transcription factors highlighted as red dots. Lists of genes for (A, B, E) highlighted in Table S2.

Loss of *PSY* receptors, whether alone (*Δpsyr1/2*) or together with sulfation (*Δpsyr1/2/Δtpst, triple knock-out*), produced only a modest, shared response (Fig. 3B). The wild type-vs-*Δpsyr1/2* and wild type-vs-*Δpsyr1/2/Δtpst* profiles were highly similar (Fig. 3B), and a direct comparison of the two mutants identified only ∼30 differentially expressed genes (Fig. 3C, S4A). Although few genes overlapped individually, only nine genes were common to wild type-vs-*Δpsyr1/2* and wild type-vs-*Δpsyr1/2/Δtpst*, all concordant in direction (Fig. 3C). This shared core looks to be non-random, converging on the same functional categories, including oxidation–reduction, protein-kinase signaling, membrane transport, and metabolic processes (12 shared Gene Ontology terms; Fig. S5A).

Four of the nine encode phenolic or cell wall enzymes: two DIR (DIRIGENT) proteins, a catechol oxidase/tyrosinase (polyphenol oxidase, PPO), and an α-L-fucosidase (cell wall glycoside hydrolase), all down-regulated (Fig. 3C). Together with a down-regulated bHLH transcription factor and a serine/threonine kinase, the two up-regulated genes encode an AP2/ERF (APETALA2/Ethylene Response Factor) transcription factor and a heavy-metal (HMA) transport/detoxification protein (Copper Transport Protein) (Fig. 3C). Notably, of these genes many are associated with stress response, including the DIR proteins, PPO, bHLH and AP2/ERF transcription factors^34–39^. The common consequence of losing PSY receptors is thus the repression of a phenylpropanoid or cell wall modules (Fig. S5B, left panel), and down-regulation of certain stress response genes (i.e., DIR proteins, bHLH, or PPO; Fig. 3C). However, the down-regulation in cell-wall associated genes (Fig. S5B, right panel) and up-regulation of the AP2/ERF transcription factors (Pp3c2_1870 and Pp3c8_7340/PpERF138) in receptor null backgrounds indicates a different signature than PSY signaling in Arabidopsis^23,24^. Although the two receptor-null lines overlapped little at the individual gene level, their differentially expressed genes converged on the same functional categories (oxidation-reduction, protein-kinase signaling, membrane transport, cell wall metabolism; Fig. S5A), suggesting that *Δpsyr1/2* and *Δpsyr1/2/Δtpst* may be functionally similar.

Because *Δpsyr1/2* and *Δpsyr1/2/Δtpst* differed only by the presence of TPST, the genes that varied between them (30 genes *Δpsyr1/2* vs *Δpsyr1/2/Δtpst*; Fig. 3C;) isolate the effect of removing sulfation in a receptor-null background, that is, TPST-dependent but PSY receptor-independent effects. As expected, expression of *TPST* itself (Pp3c16_24250) was reduced in the *Δpsyr1/2/Δtpst* mutant; which provided a quality-control confirmation rather than evidence of feedback (Fig. 3C). The remaining genes spanned phenylpropanoid/shikimate metabolism (4-coumarate-CoA ligase, DAHP synthase, histidine ammonia-lyase), photosynthesis/chloroplast functions (Photosystem II PsbM, chlorophyll a/b-binding protein, a chloroplastic Fe-superoxide dismutase), cell wall enzymes (expansin, pectate lyase, pectinesterase), abiotic-stress proteins (LEA, ferritin, catalase, a DnaJ chaperone) and several transcription factors (AP2/ERF, PLATZ) (Table S2). Of this list of genes, three candidates are shown in Fig. 3C: Pp3c17_14510 (Chloroplastic iron superoxide dismutase), Pp3c8_7340 (AP2/ERF transcription factor, PpERF138), and Pp3c13_16660 (Glutamate dehydrogenase), which are differentially expressed in at least two of the *psyr* null mutant comparisons. Given that both genotypes lack the PSY receptors, these 30 genes represent TPST-dependent and PSY receptor-independent transcriptional responses, though whether these reflect TPST activity on sulfated substrates beyond PSY remains to be determined.

In the *Δtpst* mutant, the only genotype with the PSY receptors but lacking an active sulfated ligand, the transcriptome changed strongly in a directional manner: of the 271 genes differentially expressed between wild type and *Δtpst*, the majority were down-regulated (213 down-versus 58 up-regulated; Fig. 3A, Fig. S5C). Consistent with the reduced-growth phenotype of *Δtpst* plants^27^, the repressed genes were dominated by growth-and cell wall-associated functions (Table S2, Fig. S5C): carbohydrate metabolism and cell wall modification (O-glycosyl hydrolases and pectin-lyase-fold proteins), LRR-LRKs, membrane transport, and oxidoreductase/heme-binding activities. By contrast, the upregulated gene set contained a higher proportion of transcriptional regulators: regulation of transcription and DNA-binding transcription-factor activity were among its most frequent annotations (Table S2, Fig. S5C). Among the repressed regulators were developmental transcription factors, including GRAS-family factors related to SCARECROW, SHORT-ROOT, and an SBP/SQUAMOSA factor (Fig. 3D). Notably, of the differentially regulated transcription factors, the stress-responsive WRKY family was only identified in the upregulated set (Fig. 3D). Loss of sulfation thus simultaneously represses growth and cell wall machinery and engages a transcriptional-regulator response.

The transcriptional response caused by loss of TPST depends largely on the PSY receptors. For the 262 genes differentially expressed in both, the change induced by loss of TPST (log₂ fold-change of Δ*tpst* relative to wild type; negative values indicate repression in Δ*tpst*) was strongly anti-correlated with the change induced by additionally removing the receptors (log₂ fold-change of the triple mutant relative to Δ*tpst*) (r = −0.984; Fig. 3E). In other words, genes repressed in Δ*tpst* were restored toward wild-type levels once the receptors were also removed. Consistent with this reversal, 253 of the 271 *TPST*-responsive genes behaved as PSY receptor-dependent genes (Fig. 3E). The triple mutant (*Δpsyr1/2*/*Δtpst*) confirms this dependence. The *Δtpst* mutant differed from wild type at 271 genes, whereas the *triple* differed at only 30 (Fig. 3A, S4A), suggesting that removing the receptors suppresses most of the *Δtpst* transcriptional phenotype and returns the transcriptome close to wild type.

This pattern is the opposite of what a simple activation model would predict. If sulfated PSY peptides activate the PSY receptors, removing the sulfated ligand (as is the case in the *Δtpst* mutant) and removing the receptors (*Δpsyr1/2*) would abolish the same signal and produce similar transcriptomes; instead, *Δpsyr1/2* changes only 25 genes while *Δtpst* changed 271 when compared to the wild-type transcriptome. These data provide further support for a model in which sulfated PSY restrains receptor activity: in *Δtpst*, the receptors lack their active sulfated ligand and signal constitutively, driving a growth-repressive program, while in receptor-null plants this program cannot be induced.

### Loss of PSYR kinase activity rescues gametophore development and expansion in *Δtpst*

Our data thus far suggest that PSYRs signal constitutively in the absence of a sulfated ligand, similar to what has been demonstrated in Arabidopsis^23^. Extending on this current model, we hypothesized that if unbound PSYRs continuously signal using an active kinase then PSYRs lacking kinase activity should rescue gametophore formation in *Δtpst*. Conversely, if kinase activity is activated by ligand binding, then kinase-inactive PSYRs should not rescue *Δtpst.* We directly tested this hypothesis taking advantage of the ability to rapidly and efficiently edit the genome in *P. patens*^40^. We generated a plant producing a single PSYR that is kinase inactive by introducing a mutation into the endogenous *PSYR1* locus that would result in kinase inactivation and knocking out *PSYR2*. We modeled this alteration after a well-characterized mutation (K137E) that inactivates kinase activity in Arabidopsis BAK1^41–43^. Sequence alignment of Arabidopsis BAK1 and *P. patens* PSYR1 revealed that K317 in BAK1 is conserved in PSYR1 and is located at position 843 (Fig. S6A). We designed a repair template for *P. patens PSYR1* encoding the K843E point mutation to introduce the amino acid change through HDR. We generated the point mutation and knocked out *PSYR2* in both *PSYR1-mScarlet* and *Δtpst/PSYR1-mScarlet*, allowing for visualization of the kinase-dead receptor (Fig. S6B, Table S1). From this point on, *psyr1-K843E-mScarlet*/*Δpsyr2* and *Δtpst/psyr1-K843E*-*mScarlet*/*Δpsyr2* are referred to as *psyr1-KD* and *Δtpst/psyr1-KD.* Both *psyr1-KD* and *Δtpst/psyr1-KD* formed expanded gametophores and displayed normal senescence (Fig. 4A). Similar to *Δpsyr1/2*, *psyr1-KD* was 26% larger than wild type (Fig. 4B). However, unlike *Δpsyr1/2/Δtpst*, *Δtpst/psyr1-KD* was similar in size to wild type (Fig. 4B). Both *psyr1-KD* and *Δtpst/psyr1-KD* showed normal first cell divisions during gametophore formation and showed normal bud morphology compared to wild type (Fig. 4C). Single gametophores of *psyr1-KD* and *Δtpst/psyr1-KD* were similar to wild type (Fig. 4D), indicating that gametophore development was undisturbed by the *psyr1-KD* mutation, and that *psyr1-KD* rescues *Δtpst*.

**Figure 4.**
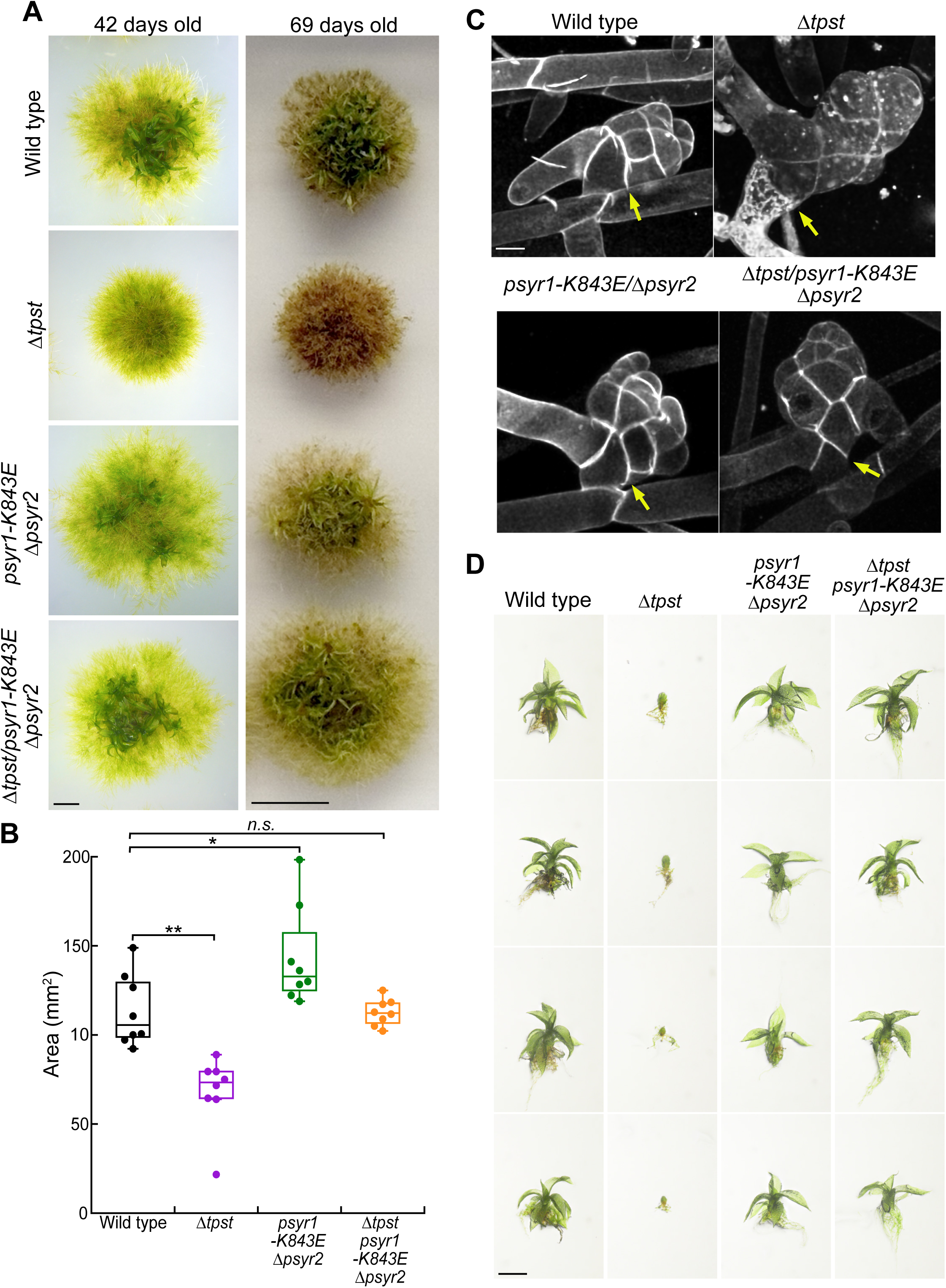
PSYR1 kinase inactivation rescues *Δtpst*. All depicted lines have endogenously tagged *PSYR1-mScarlet*. A) Representative brightfield images of 42-day-old wild-type, *Δtpst, psyr1-K843E/Δpsyr2*, and *Δtpst/psyr1-K843E/Δpsyr2* plants regenerated from protoplasts. Scale bar, 0.2 cm. Images on the right depict the same plants at 69-days-old. Scale bar, 0.5 cm. B) Quantification of plant area of 42-day-old plants. Significant differences determined by a Kruskal-Wallis test. **Wild type vs *Δtpst*, p=0.0016; *Wild type vs *psyr1-K843E*/*Δpsyr2*, p=0.03569. n.s., not significant. C) Representative maximum intensity projections of confocal z-stacks from propidium-iodide-stained buds. Yellow arrow denotes the first division in the bud initial. Scale bar, 20 µm. D) Representative extended depth of focus brightfield images of single gametophores dissected from 4-week-old ground tissue. Scale bar, 0.5 mm.

To assess if the psyr1-K843E-mScarlet localizes to the plasma membrane, we imaged individual gametophores. We found that both *psyr1-KD* and *Δtpst/psyr1-KD* had mScarlet fluorescence levels that resembled PSYR1-mScarlet (Fig. 5A). Compared to *Δtpst/PSYR1-*mScarlet, there was noticeably less PSYR1-mScarlet on the plasma membrane in *Δtpst/psyr1-KD* (Fig. 5A), indicating that the *psyr1-KD* mutation results in decreased levels of the receptor on the membrane in the *Δtpst background*. Furthermore, the morphology of *Δtpst/psyr1-KD* cells in the gametophore was oblong and regular, similar to the wild type and differing from the more compact and irregularly shaped cells in a *Δtpst* aborted gametophore. We found that PSYR1-mScarlet fluorescence levels in *Δtpst/*PSYR1-mScarlet were 2.4 times greater compared to PSYR1-mScarlet and *psyr1-KD*, and 2.2 times greater than *Δtpst/psyr1-KD* (p < 0.001) (Fig. 5B). Neither *psyr1-KD* nor *Δtpst/psyr1-KD* exhibited levels of PSYR1-mScarlet fluorescence that were different from *PSYR1-mScarlet* (Fig. 5B). Because the kinase-dead receptor no longer accumulates in the *Δtpst* background despite being present, these data indicate that receptor accumulation depends on kinase activity rather than on loss of peptide sulfation. Decoupling the receptor’s presence at the membrane from its activity indicates that the *Δtpst* phenotype is driven by kinase activity rather than by the amount of membrane-associated receptor.

**Figure 5.**
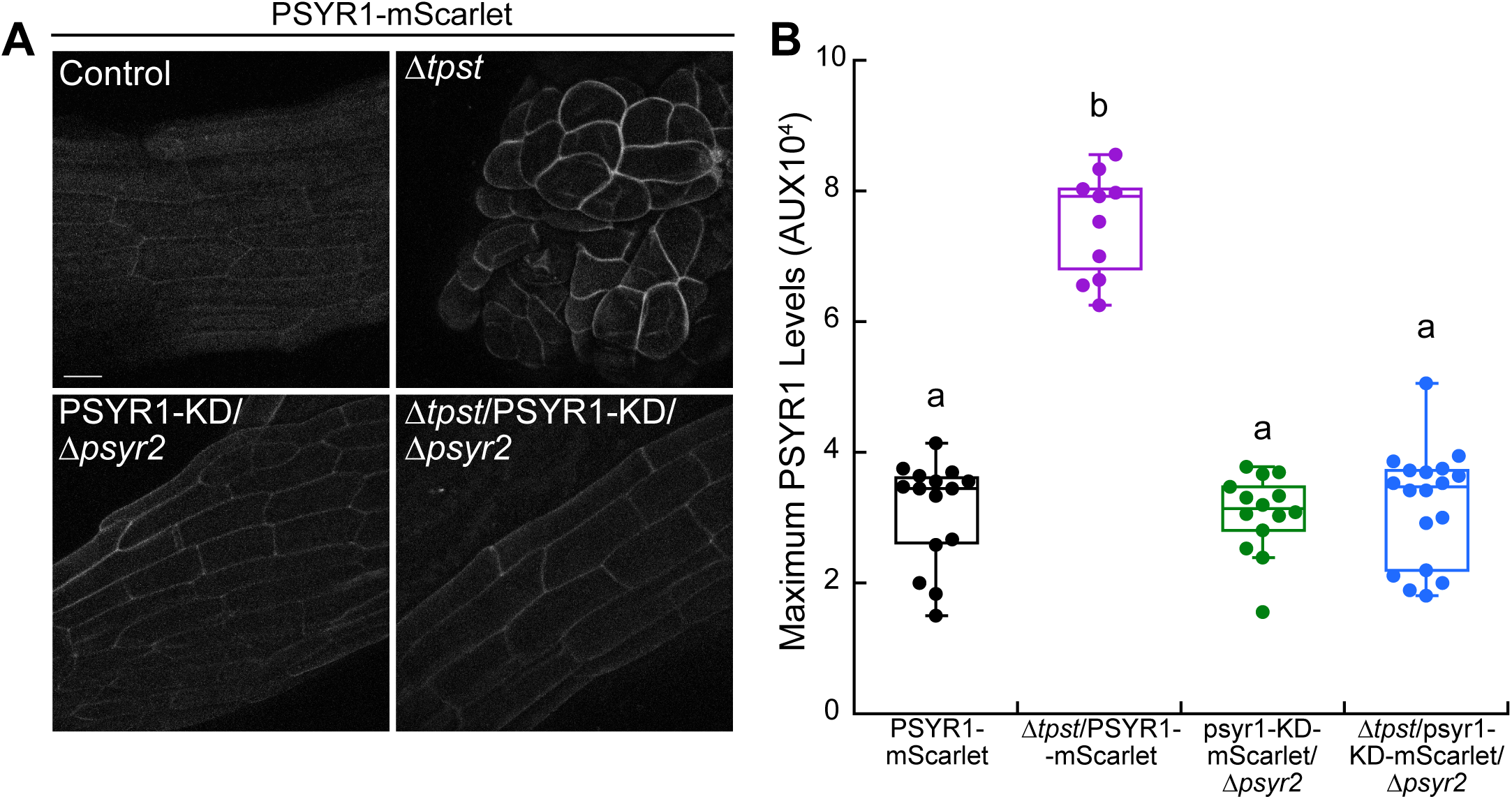
PSYR1 kinase inactivation alters membrane-associated PSYR1 levels in *Δtpst.* A) Maximum intensity projections from confocal z-stacks of phyllids in expanded gametophores, or large aborted buds (*Δtpst*), dissected from approximately 4-week-old ground tissue. Scale bar, 20 µm. B) Quantification of PSYR1-mScarlet fluorescence intensity from maximum intensity projection images. Significant differences determined by a Kruskal-Wallis test with a Dunn’s post hoc test and Bonferroni correction (α=0.05) are indicated by different letters.

Because *Δpsyr1/2/Δ*tpst, but not *Δtpst/psyr1-KD* exhibited increased 2D growth compared to wild type, we sought to understand how the PSYR kinase-dead mutant differed from the null mutant. To do this, we examined the expression levels of *PSY, PSYR1,* and *PSYR2* in both the null mutant and kinase-inactive lines. In the null receptor mutant lines, the expression levels of each selected gene in *Δpsyr1/*2 and *Δpsyr1/2/Δtpst*, was not significant (Fig. 6A). Interestingly, both *psyr1-KD* and *Δtpst/psyr1-KD* had opposing expression levels of *PSY, PSYR1,* and *PSYR2* compared to the *Δpsyr1/2* and *Δpsyr1/2/Δtpst*. The *psyr1-KD* and *Δtpst/psyr1-KD* mutants showed a dramatic 14.1- and 19.2-fold upregulation of *PSY* expression, respectively (Fig. 6B). Levels of *PSYR1* and *PSYR2* expression were less dramatic than *PSY* but still significant, with *psyr1-KD* and *Δtpst/psyr1-KD* exhibiting a 2.8- and 3.6-fold increase in *PSYR1* and a 2.7- and 3.1-fold increase in *PSYR2* expression, respectively. Despite the upregulation in *PSYR1,* both *psyr1-KD* and *Δtpst/psyr1-KD* did not exhibit any difference in levels of PSYR1-mScarlet fluorescence on the membrane (Fig. 5B), suggesting that transcript and protein levels are uncoupled. In *psyr1-KD* and *Δtpst/psyr1-KD*, there were no significant differences in *PSY, PSYR1,* and *PSYR2* expression levels. However, expression of *PSY*, *PSYR1*, and *PSYR2* was significantly different between the kinase-dead mutants and *Δtpst* (Fig. 6B). In contrast, there were no differences in expression levels of both *PSYR1 and PSYR2* between wild type, *Δtpst,* and the *psyr* receptor null mutants (Fig. 6A). Additionally, there was no significant difference in *PSY* levels between wild type and the *psyr* receptor null mutants, although *PSY* was downregulated in *Δtpst.* These data indicate that the transcriptional effects of a *psyr1* and *psyr2* null mutation are less pronounced than those in the *psyr1-KD* mutant. Because the null receptor and the kinase-dead receptor mutants exhibit different expression levels of *PSY, PSYR1,* and *PSYR2*, these data suggest that there is a fundamental difference between the absence of the receptors from the membrane and inactivation of the receptors for controlling expression of *PSY* and its receptors.

**Figure 6.**
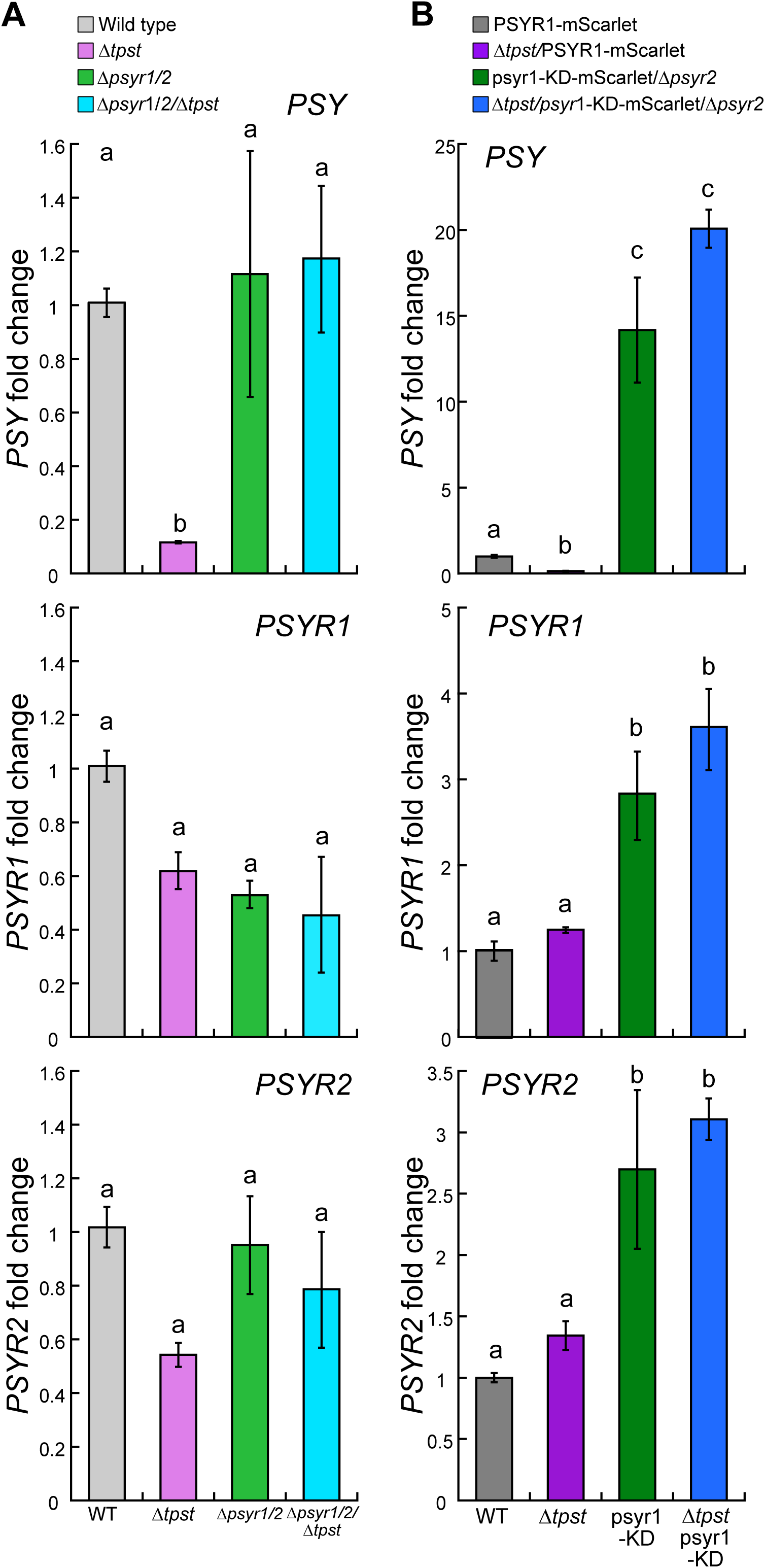
PSYR1 kinase inactivation upregulates *PSY* and *PSYR* expression. A) Expression level of *PSY*, *PSYR1* and *PSYR2* in approximately 4-week-old ground tissue of *psyr* null (A) or *psyr1-*KD (B) mutants. Different letters indicate significant differences determined by a one-way ANOVA with Tukey’s multiple comparisons (α=0.05).

### Overexpression of PSYRs phenocopy Δtpst

Because *Δtpst* has high levels of PSYRs on the plasma membrane, and removal of the receptors rescues *Δtpst*, we predicted that PSYR overexpression would phenocopy *Δtpst*. To test this, we generated overexpression lines for either PSYR1-mStayGold or PSYR2-mStayGold in the *PSYR1-mScarlet* background to allow for differentiation between the endogenous PSYR1 receptor and the overexpressed PSYR1 or PSYR2 receptor. We used CRISPR-mediated HDR to insert the *PSYR1* or *PSYR2* coding sequence fused to mStayGold into a non-essential locus^44^ in the *P. patens* genome. The constructs were driven by the rice actin promoter for constitutive expression^45,46^. Overexpression lines were identified by screening for mStayGold fluorescence in protonemata (Fig. S7B, Table S1), a developmental stage where PSYRs are expressed at very low levels. Two PSYR1 overexpression lines, *PSYR1-mScarlet/ACT1::PSYR1-mStayGold* #22 and #23 were identified and hereafter referred to as *PSYR1-OE* #22 and *PSYR1-OE* #23, respectively. One PSYR2 overexpression line, *PSYR1-mScarlet/ACT1::PSYR2-mStayGold* #9 (hereafter referred to as *PSYR2-OE*) was also isolated. Both *PSYR1-OE* #22 and *PSYR2-OE* plants were round, dense, and unable to form expanded gametophores, very similar to *Δtpst* (Fig. 7A). However, *PSYR1-OE* #23 formed gametophores but exhibited fewer protonemal protrusions from the body of the plant compared to wild type, suggesting an intermediate phenotype (Fig. 7A). Interestingly, *PSYR1-OE* #22 displayed accelerated senescence, turning brown faster than *Δtpst* (Fig. 7A).

**Figure 7.**
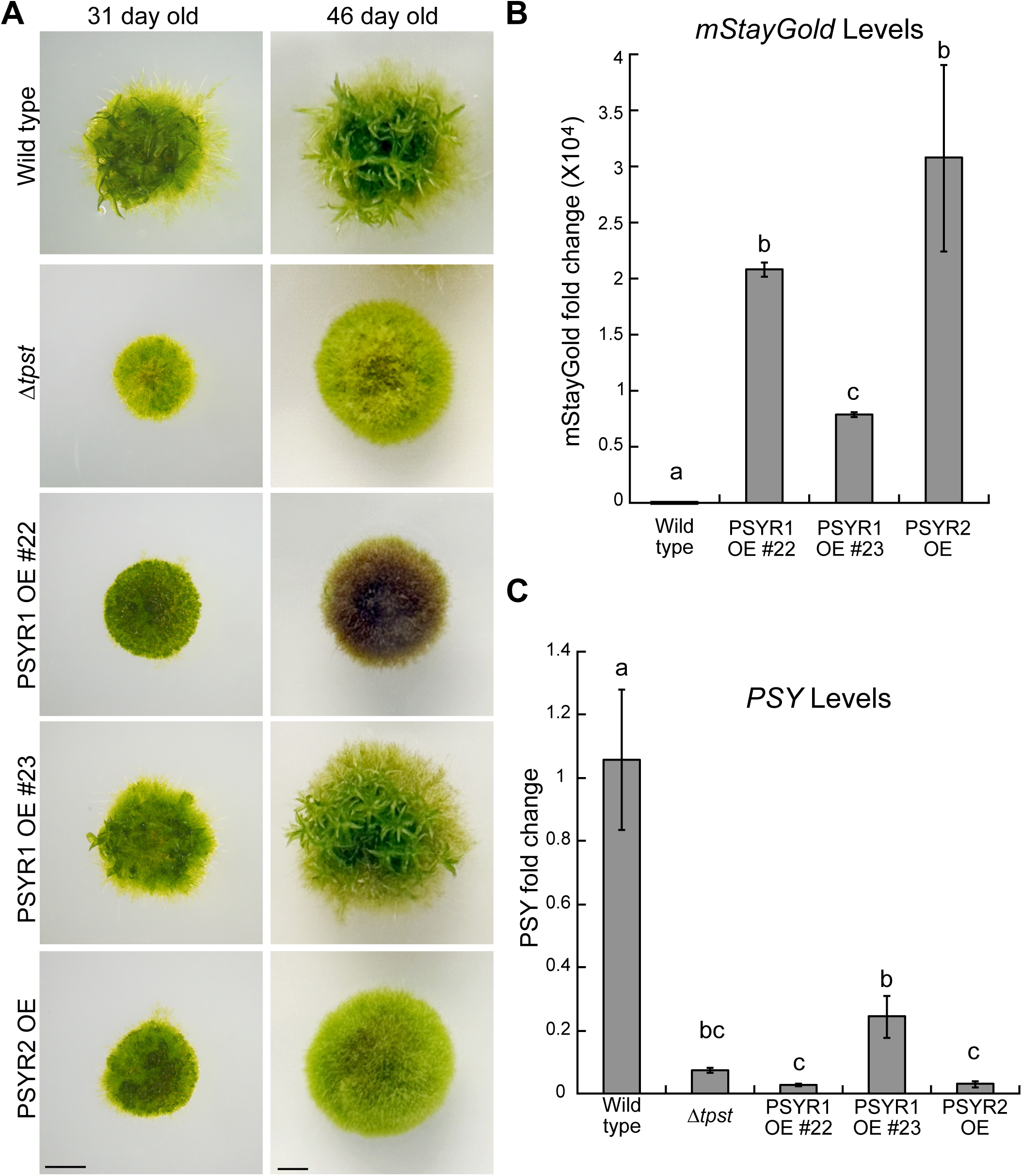
PSYR overexpression phenocopies *Δtpst*. A) Representative images of 31-day-old plants regenerated from protoplasts. Images on the right depict the same plants at 46-days-old. Scale bar, 0.2 cm. B) *PSY* and (C) *mStayGold* expression levels in approximately 3-week-old ground tissue. Significant differences determined by a one-way ANOVA with Tukey’s multiple comparisons (α=0.05) are indicated by different letters.

To correlate phenotype with expression levels, we used RT-qPCR to measure the transcript levels of mStayGold in each line. Both *PSYR1-OE* #22 and *PSYR2-OE* exhibit levels of mStayGold expression that are not statistically different from each other (Fig. 7B), but are significantly higher than *PSYR1-OE* #23 (Fig. 7B). These data suggest that levels of *PSYR* correlate with the severity of the *Δtpst*-like phenotype. Interestingly, *PSYR1-OE* #22 displays accelerated early senescence compared to *PSYR2-OE* despite similar mStayGold levels, raising the possibility that PSYR1 may contribute more to senescence as compared to PSYR2. Additionally, all three overexpression lines and *Δtpst/PSYR1-mScarlet* have decreased levels of *PSY* compared to wild type (Fig. 7C). Taken together, our data suggest that overexpression of the PSYRs recapitulates the *Δtpst* phenotype, both morphologically and transcriptionally.

We used CRISPR-mediated genome editing to introduce the corresponding kinase inactivating mutations (PSYR1-K843E and PSYR2-K841E) into the *PSYR1-OE #22* and *PSYR2-OE* lines, yielding *PSYR1-OE #22/psyr1-K843E* and *PSYR2-OE/psyr2-K841E* mutant lines (hereafter referred to as *psyr1-KD-OE* and *psyr2-KD-OE*) (Fig. S8A-B, Table S1). We found that the kinase-dead mutations rescued gametophore formation in both *psyr1-OE #22* and *psyr2-OE* (Fig. S8C), and *psyr1-KD-OE* did not show accelerated senescence (Fig. S8D). Furthermore, both *psyr1-KD-OE* and *psyr2-KD-OE* show increased *PSY* expression that is approximately 2.5 times greater than wild type (p=0.011 and 0.012, respectively) and approximately 37 times greater than *Δtpst* (p<0.0001) (Fig. S8E). These data confirm that the overexpression phenotypes are due to enhanced levels of PSYRs with active kinases, further supporting that increased kinase activity, rather than the physical presence of the receptor, phenocopies *Δtpst*. Moreover, since both *psyr1-KD-OE* and *psyr2-KD-OE* have similar phenotypes, these data indicate that either a PSYR1 or PSYR2 kinase-inactive mutation is sufficient to rescue gametophore expansion, suggesting that PSYR1 and PSYR2 kinase activity behave similarly as negative regulators of growth.

## Discussion

Here, we provide multiple lines of evidence that PSY’s roles in promoting cellular proliferation and expansion are evolutionarily conserved. We identified and characterized the receptors in *P. patens* (PSYR1/Pp3c24_159505 and PSYR2/Pp3c8_17530) that are necessary for PSY signaling and found that *psyr* null mutants are insensitive to PSY and have enhanced plant growth. Although a previous study reported five putative *P. patens* PSY receptor orthologs^26^, three of these candidates are unlikely to be PSY receptors based on the number of LRRs and lack of conserved sulfotyrosine binding residues (referenced in Fig. S1). The other two candidates correspond to the orthologs verified in our study (Pp3c24_159505 and Pp3c8_17530). Using endogenous tagging of the receptors, we showed that levels of membrane-associated receptors are critical for fine-tuning cell expansion, and overexpression of the receptors inhibited growth. Furthermore, our data support a model whereby receptor kinase activity is inversely related to cell expansion.

PSY’s role in enhancing cellular proliferation in the root apical meristem and diffuse expansion in the root elongation zone in rice and Arabidopsis is supported by exogenous treatment or overexpression of PSY1 inducing root elongation^9,14,30,31^. In *P. patens* ground tissue, which is comprised of densely populated plants, exogenous PpPSY1 promoted bud and gametophore formation. The increased number of buds and gametophores parallels the enhanced root growth in angiosperms since bud and gametophore formation relies on the transition to 3D growth by the establishment of a proliferative meristematic cell and diffuse expansion of the gametophore leaflets. In contrast, *Δpsyr1/2* was insensitive to PSY1, providing evidence that these receptors are necessary for PSY recognition. The PSY-insensitive *Δpsyr1/*2 mutant had more buds and gametophores compared to wild type under control conditions, suggesting that *psyr* null mutants display a constitutive PSY response.

We found that bud and gametophore formation differed when plants were grown from individual protoplasts and isolated from one another. In this case, *psyr* null mutants exhibited increased 2D growth and a reduced number of expanded gametophores, suggesting that the transition from 2D to 3D growth may depend on protonemal tissue density. In comparison, *Δtpst* showed decreased protonemal and gametophore expansion regardless of tissue density. Furthermore, in the absence of TPST, both PSYR1 and PSYR2 are significantly enriched on the membrane despite no significant difference in *PSYR1* and *PSYR2* transcript levels. These data suggest that PSYRs negatively regulate cell expansion.

Thus, in the absence of the receptors with or without TPST, we would expect an increase in cell expansion, which is what we observed. Interestingly, the increased surface area covered by the *psyr* null mutants resembles observations in Arabidopsis where the *psyr1,2,3* mutant has longer roots^23^. However, the *tpst/psyr1,2,3* quadruple mutant did not exhibit the enhanced root elongation of *psyr1,2,3* but instead was comparable to PSY5-treated *tpst-1* roots, indicating that removing the PSY receptors only partially relieves the *tpst* defect in Arabidopsis. By contrast, in *P. patens* the enhanced plant area in *Δpsyr1/2* and *Δpsyr1/2/Δtpst* is indistinguishable. This suggests that in *P. patens,* a *psyr* null mutant is fully epistatic to *Δtpst*, while only partially epistatic in Arabidopsis. To date, *P. patens* has only one functional class of tyrosine-sulfated peptides, PSY^25–27^. The previously identified PpPSK peptide^25^ lacks the conserved DY residue required for tyrosine-sulfation, suggesting it is not sulfated^27^. However, Arabidopsis has multiple classes of tyrosine-sulfated peptides (PSY, PSK, RGF, CIF), each perceived by distinct receptors, which may explain why removing the PSY receptors alone is only partially epistatic in Arabidopsis. Moreover, root elongation is undoubtedly more complex than protonemal tissue expansion and likely requires additional signaling modules.

The expanded protonemal plant area in the *psyr* null mutants, *Δpsyr1/2* and *Δpsyr1/2/Δtpst*, is consistent with the observed differential expression of cell wall-associated genes (Fig. S5B, Table S2), including a putative pectin methylesterase inhibitor (Pp3c10_1100; PMEI), alpha-L-fucosidase (Pp3c18_1550, important for xyloglucan synthesis), and two DIR-like proteins (Pp3c6_6545, Pp3c5_22560), all of which are down-regulated and can influence cell wall stiffness and reinforcement^47–49^. Notably, in *Arabidopsis,* loss of the PSY receptors (*tri-1/atpsyr1,2,3*) instead induces wall-loosening enzymes, with expansins and xyloglucan endotransglucosylases/hydrolases predominantly up-regulated (16 up, 7 down; Fig. S5B, Table S2), consistent with the up-regulation of cell wall-remodeling genes reported as the central molecular finding of^24^. Reducing genes associated with cell-wall stiffening/reinforcement in moss and increasing cell-wall loosening enzymes in *Arabidopsis* represent two distinct routes toward the same predicted outcome, a more extensible cell wall. This shared outcome is consistent with the enhanced growth of both receptor-null mutants, which form larger plants in moss (Fig. 2A,B) and longer roots in *Arabidopsis*^23,24^. When comparing *P. patens* DEGs to Arabidopsis, only 49% (1,104 of 2,237) in our study had an assignable Arabidopsis ortholog (Table S2; Ensembl Plants; mean 39% identity). Therefore, although the two systems share a small number of possible orthologous genes, they converge on a common physiological outcome through opposite molecular strategies (Fig. S5B, Table S2). Loss of the PSY receptors enhances growth in both moss and *Arabidopsis*, and in both cases, this is accompanied by transcriptional changes predicted to modulate cell-wall structure. The two lineages reach this state; however, the results suggest through different gene sets and targeted pathways (Fig.5SB; Table S2). These observations suggest that the growth-repressive function of the PSY receptors may be ancient, whereas the specific downstream targets have diverged.

Furthermore, the *P. patens psyr* null mutants showed upregulation of the AP2/ERF transcription factors (Pp3c2_1870 and Pp3c8_7340/PpERF138, Fig. 3C), which differs from PSY signaling in Arabidopsis^23^. This suggests that although the PSY signaling pathway and its growth output are conserved, the exact downstream transcriptional targets differ between the two lineages. In addition, the *psyr* null mutants showed downregulation of certain stress-response genes (Ex. Glutamate dehydrogenase, FE-SOD, DIR-proteins, and PPO; Fig. 3C). However, despite the prominent expansion phenotype and these transcriptional differences, *Δpsyr1/2* and *Δpsyr1/2/Δtpst* transcriptomes strikingly only differed from wild type by 25 and 30 genes, respectively. Thus, in *P. patens* loss of the PSY receptors produces a small, but focused transcriptional response.

On the other hand, in *Δtpst* and therefore in the absence of sulfated peptides, there were 271 genes exhibiting at least two-fold differential expression compared to wild type. Gene ontology analysis revealed that loss of tyrosine sulfation represses cell wall and growth machinery, while simultaneously engaging a transcriptional regulator response consistent with enhanced stress response and decreased developmental transitions. Specifically, a WRKY family transcription factor is upregulated only in *Δtpst* (Fig. 3D). WRKY transcription factors play important roles in biotic and abiotic stress, as well as development, including senescence^50,51^. GRAS-family transcription factors related to SHORT-ROOT and SCARECROW are downregulated in *Δtpst*, and they are important for regulating periclinal divisions during leaf blade and mid-vein formation in the gametophores of *P.* patens^52,53^. Additionally, a SQUAMOSA PROMOTER BINDING-LIKE PROTEIN (SPL) was upregulated in *Δtpst*. In Arabidopsis, SPL proteins promote the transition from juvenile to adult tissue fate^54,55^. Together, these downregulated genes are consistent with *Δtpst*’s inability to form expanded phyllids.

Of note, the majority (253 of 271) of TPST-responsive genes are PSY receptor-dependent genes, supporting a model in which PSYRs signal constitutively to repress growth in the absence of a sulfated ligand, like in *Δtpst*. However, binding of PSY restrains receptor activity, promoting growth, suggesting that *psyr* null mutant lines cannot induce the growth-repression program, and further supports that *psyr* null mutants exhibit a constitutive PSY response. Transcriptional profiles for stress-responsive genes are inversely correlated (Fig. 3D), with upregulation in *Δtpst* and downregulation in *psyr* null mutants. This observation suggests that in addition to repressed growth in the absence of ligand, PSYRs are also promoting the stress response, aligning with the “ligand-deprivation-dependent activation system” used to describe PSY signaling in Arabidopsis^23^.

Because growth and expansion are promoted in the presence of the PSY peptide, we sought to determine whether ligand binding results in kinase activation in accordance with the prevailing mechanism for LRR-RLKs^56^. If peptide binding were to lead to kinase activation, then we would predict that a kinase-inactive mutant would not rescue *Δtpst*. However, the kinase-inactive mutant, *psyr1-K843E/Δpsyr2* (*psyr1-KD*), fully rescued *Δtpst.* Notably, the *psyr1-*KD mutation in *Δtpst* decreases PSYR1-mScarlet levels on the membrane to the same intensity as wild type, suggesting that PSYR1 kinase activity modulates PSYR membrane association, with active kinase leading to increased levels and inactive kinase leading to decreased levels. Thus, PSY binding likely serves to inactivate, rather than activate, the PSYR kinase, which results in reduced membrane-associated PSYR. To our knowledge, this is the first example of an LRR-RLK with an active kinase domain in the absence of a ligand that appears to be inactivated in conjunction with ligand binding. In fact, a kinase inactivation model is the opposite of the proposed signaling mechanism for the PSK, in which the kinase activity of its receptor, PSKR1, is required for PSK signaling, and PSKR1 is phosphorylated upon PSK binding^57,58^.

The *PSY* transcriptional profile further supports that kinase inactivation is critical for transitioning to growth. *PSY*, which is growth-promoting, was significantly upregulated in kinase-inactive receptor mutants but not in *null* receptor mutants. Additionally, overexpression of either PSYR1 or PSYR2 resulted in decreased levels of *PSY*, while overexpression of kinase-inactive PSYR1 or PSYR2 enhanced *PSY* levels. Of note, the PSYR-overexpression lines exhibiting the highest levels of *mStayGold* expression (PSYR1-OE #22 and PSYR2-OE) showed the lowest levels of *PSY* expression. Taken together, our data suggest that the amount of PSYR with an active kinase domain predicts the extent to which *PSY* expression is inhibited. If the proportion of active to inactive PSYR kinase serves to tune *PSY* transcript levels, then the absence of PSYR, as is the case in the *psyr* null mutants, should have no effect on *PSY* levels, which is what we observed. Similarly, the muted transcriptional response in the *psyr* null transcriptomes could be attributed to the absence of the kinase. Finally, the varying phenotype between *PSYR1-OE #22, PSYR2-OE,* and *PSYR1-OE #23* suggests that the amount of PSYRs is directly related to the levels of growth inhibition and, therefore, phenotypic severity.

By investigating PSYR function in *P. patens*, a relatively simple plant with rapid genome editing capabilities and fewer LRR-RLKs and sulfated peptide classes compared to angiosperms, our data has provided key mechanistic insights into this complex signaling pathway. We have found that PSYRs localize to the plasma membrane and serve as a growth inhibition signal via an active kinase, which results in reduced levels of *PSY* (Fig. 8). In the absence of ligand, kinase active PSYRs continually repress growth and accumulate on the membrane. However, binding of PSY serves to inactivate the receptors by turning the kinase domain off, relieving the growth inhibition signal and resulting in reduced levels of membrane-associated PSYR. In this model, fine-tuning growth relies on negative feedback between membrane-associated PSYR kinase activity, which is growth repressive, and transcriptional control of the *PSY* ligand, which reduces PSYR kinase activity.

**Figure 8.**
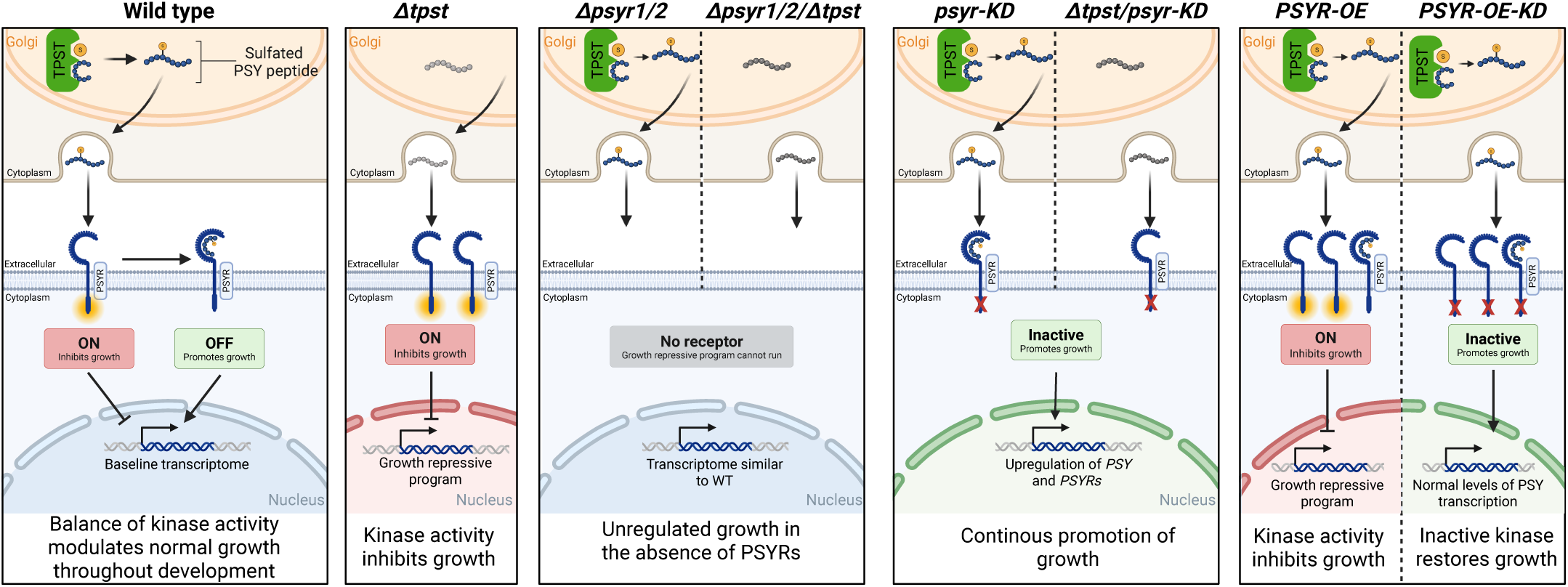
Diagram representing how each mutation affects PSY signaling and growth. In wild type, sulfated PSY produced by TPST binds to PSYR, which turns off the PSYR kinase and results in the basal transcriptome modulating normal growth throughout development. When sulfated PSY is absent, as is the case in the absence of *TPST* (*Δtpst*), unbound PSYR remains active, triggering a growth-repressive transcriptional program. In the receptor null mutants (either *Δpsyr1/2* or *Δpsyr1/2/Δtpst*), the receptors are absent and the growth repressive program is not triggered, resulting in ligand-independent unregulated growth with transcriptomes similar to wild type. In contrast to the absence of PSYR, kinase-dead PSYR induces transcriptional changes such as upregulating *PSY*, *PSYR1*, and *PSYR2* expression, and promotes growth. Overexpression of PSYRs leads to growth inhibition due to more active membrane-associated kinase and less PSY peptide expression, which can be relieved by inactivating the kinase. Figure made with BioRender.com

## Materials and Methods

### Tissue propagation

*P. patens* lines were propagated weekly by moderate homogenization in water and then plated onto PpNH4 (1.03 mM MgSO4, 1.86 mM KH_2_PO_4_, 3.3 mM Ca(NO_3_)_2_, 2.7 mM (NH_4_)_2_-tartrate, 45 μM FeSO_4_, 9.93 μM H_3_BO_3_, 220 nM CuSO_4_, 1.966 μM MnCl_2_, 231 nM CoCl_2_, 191 nM ZnSO_4_, 169 nM KI, and 103 nM Na_2_MoO_4_, containing 0.7% agar) plates covered with cellophane^44^. Tissue was grown in Percival growth chambers at room temperature under 85 µmol_photons_/m^2^s light with long-day (16/8) conditions. Peptide media was created by supplementing PpNH4 with 1 µM PpPSY1.

### Identification of putative P. patens PSY Receptor orthologs

The validation of PSY receptor orthologs was done using NCBI BlastP against the NCBI database: *A. thaliana* (taxid: 3702) and *P. patens* (taxid:3218). The full-length protein sequences for *Arabidopsis thaliana* PSYR1 (At1g17230), PSYR2 (At2g33170), PSYR3 (At5g63930), and RGFR1 (At4g26540) were acquired from the TAIR repository (https://www.arabidopsis.org/). These sequences served as queries for BlastP analyses and sequence alignments. The full-length protein sequences for *P. patens* PSY receptor candidates (Pp3c24_15950, Pp3c8_17530) and putative *O. sativa* PSY receptors LOC_Os07g05740, LOC_Os04g42700) were acquired from Phytozome (https://phytozome-next.jgi.doe.gov/^59^) and used for reciprocal BlastP analyses and sequence alignments.

### Plasmid and oligonucleotide construction

CRISPR targeting plasmids were created using a single-stranded oligo containing the protospacer sequence and the pMH-Cas9-sgRNA plasmid backbone^40^. All oligos are shown in Table S3. All plasmids were assembled using Hi-Fi DNA Assembly Mastermix (NEB). All HDR plasmids and oligos were designed as previously described^40^. The PSYR1-mScarlet and PSYR2-mScarlet vectors were constructed using a pGEM backbone^40^. The PSYR1-mStayGold and PSYR2-mStayGold overexpression plasmids were constructed by PCR-linearizing the pTKUbi vector and replacing the maize ubiquitin promoter with the rice actin 1 promoter^44^. All plasmids were sequence verified.

### Generation of knockout and point mutation lines

Approximately one-week-old *P. patens* tissue was digested to generate protoplasts, which were transformed using a PEG-mediated transformation protocol^40^. To generate knockouts, 15 µg of the CRISPR plasmid was used. For HDR transformations using annealed oligonucleotides (*psyr1-K843E, psyr2-K841E)*, 15 µg of the CRISPR plasmid and 10 µL of annealed oligonucleotides were used. The *tpst* null mutant in *Δpsyr1/2/Δtpst* was created using an oligo encoding a stop codon. All oligo sequences are listed in Table S3. Details for each generated line are listed in Table S1.

After transformation, protoplasts were recovered and placed on selection by one of two methods: 1) Protoplasts were plated on cellophane-covered PRMB media (PpNH4 medium supplemented with 8.5% mannitol and 10 mM CaCl_2_) for 4 days. The cellophane, which contains the protoplasts, was moved to selection media (PpNH4 media containing 15 µg ml^-1^ hygromycin) for 1 week, and then moved to PpNH4^40^. 2) Protoplasts were left in a suspension of 3mL of liquid plating media (4 mM KNO_3_, 2 mM KH_2_PO_4_, 1 mM Ca(NO_3_)_2_, 89 µM Fe citrate, 300 µM MgSO_4_, 9.93 µM H_3_BO_3_, 220 nM CuSO_4_, 1.966 µM MnCl_2_, 231 nM CoCl_2_, 191 nM ZnSO_4_, 169 nM KI, 103 nM Na_2_MoO_4_, supplemented with 8.5% mannitol and 9.8 mM CaCl_2_). After 4 days, protoplasts were centrifuged for 7 min at 250 × *g*, room temperature, and plated onto cellophane-covered PRMB selection media (PRMB media containing 15 µg ml^-1^ hygromycin) for 1 week. After 1 week, the cellophane was moved to PpNH4.

Surviving protoplasts have taken up the CRISPR plasmid. After 1-2 weeks, protoplast-regenerated plants were individually picked to fresh PpNH4 plates without cellophane, before being genotyped around 2 weeks later.

### Genotyping

Genomic DNA was extracted from *P. patens* tissue^40^. Competition PCR was used to genotype plants for null and point mutations and plants with endogenous tagging were genotyped using PCR primers upstream and downstream of the 5’ and 3’ flanking sequences to screen for plants with an insertion^40^. For genotyping plants with an overexpression construct, a forward primer in the receptor coding sequence and a reverse primer in the mStayGold sequence were used to confirm integration of the HDR construct. Positive plants were screened for mStayGold overexpression using confocal microscopy. All PCR products were sequenced using Sanger sequencing. All genotyping primers are listed in Table S3.

### Laser scanning confocal microscopy

Agar pads were made on glass microscope slides and covered with Hoagland’s medium (4 mM KNO_3_, 2 mM KH_2_PO_4_, 1 mM Ca(NO_3_)_2_, 89 µM Fe citrate, 300 µM MgSO_4_, 9.93 µM H_3_BO_3_, 220 nM CuSO_4_, 1.966 µM MnCl_2_, 231 nM CoCl_2_, 191 nM ZnSO_4_, 169 nM KI, 103 nM Na_2_MoO_4_)^44^. To screen for fluorescence, sections of protonemal tissue were cut from the cellophane and transferred to the pad, and then covered with a glass coverslip. For imaging gametophores, single gametophores were picked from ground tissue and placed directly onto Hoagland’s-covered agar pad. All samples were sealed with VALAP (1:1:1 parts of Vaseline, lanoline, and paraffin).

For propidium iodide imaging, all lines except *Δtpst* were grown on PpNO_3_ (1.03 mM MgSO4, 1.86 mM KH_2_PO_4_, 3.3 mM Ca(NO_3_)_2_, 45 μM FeSO_4_, 9.93 μM H_3_BO_3_, 220 nM CuSO_4_, 1.966 μM MnCl_2_, 231 nM CoCl_2_, 191 nM ZnSO_4_, 169 nM KI, and 103 nM Na_2_MoO_4_, containing 0.7% agar). *Δ*tpst was grown on PpNH4. Sections of approximately 1.5-week-old ground tissue was cut from the cellophane and transferred to an agar pad covered with 10-20 µg/mL of propidium iodide in 0.5x Hoagland’s and incubated in the dark for 10 minutes.

Images were collected using a 0.75 NA 20X objective (Nikon). Laser illumination was set to 488 nm for mStayGold (laser power: 2%, Gain: 60; PMT offset: 28) and 561 nm for mScarlet (laser power: 1-2%; Gain: 35-60; PMT offset: 20-24). Emission filters were 525/50 nm for mStayGold and 595/50 nm for mScarlet. For propidium iodide, laser illumination was set to 561 nm (laser power: 1%; Gain: 85; PMT offset: 24).

### RNA expression analysis

Total RNA was isolated from approximately 3-4-week-old ground tissue regenerated from protoplasts using a RNeasy Plant mini prep kit (Qiagen). cDNA was prepared using 250-500 µg of total RNA using Superscript III reverse transcriptase (Invitrogen). RT-qPCR was performed using iTaq universal SYBR Green Supermix (Bio-Rad) with three biological replicates and two technical replicates, using *elongation factor 1-alpha* (Pp3c2_6770) for normalization. All primers are listed in Table S3.

### Brightfield imaging and morphometric analysis

Protoplast preparation was performed using a standard protocol using approximately one-week-old ground tissue^40^. Protoplasts were plated onto cellophane-covered PRMB media. After 4 days, the protoplasts on cellophane were moved to PpNH4 media.

All brightfield images were collected using a stereomicroscope (Nikon SMZ25) equipped with a CCD color camera (Nikon digital DS-Fi2) with a 1X objective. Acquisition settings were held at the same settings for each image. To quantify plant area, the blue channel was separated from each RGB image, which encapsulates the area of the plant. The image was inverted and thresholded. The area of the thresholded image was measured in Fiji. To image gametophores, single gametophores were picked from approximately 4-week-old ground tissue grown on PpNH4. Dissected gametophores were placed on a new plate of PpNH4 and laid on their side for imaging. Extended depth of focus images were processed using NIS elements software (Nikon).

### Fluorescence intensity quantification

Maximum intensity projection from confocal Z-stacks of single dissected gametophores were acquired. Acquisition settings were held constant. At least 10 images were collected per line. A box of a given size (140.87 µm x 74.16 µm) was placed over the image. We plotted the profile of the box in Fiji and used a macro written in R to measure the area under the curve. Quantification was performed using R version 4.4.1^60^.

### Statistical Analyses

A non-parametric approach was taken to analyze significant differences. For differences in plant area between mutant lines and wild type, and the formation of 3D structures in control versus peptide-treated conditions, statistical significance was determined using a Kruskal-Wallis test with a chi-squared approximation. For differences in fluorescent signal, a Kruskal-Wallis test followed by a post-hoc Dunn’s test was used to determine significance. A Bonferroni correction was applied to the p-values resulting from Dunn’s test to control for multiple comparisons. A significance level of α = 0.05 was used to identify groups that were statistically different.

For RT-qPCR expression data, a one-way ANOVA with a post-hoc Tukey HSD test was used to determine significant groups based on the ΔCt values and reported on graphs of relative of expression. For all multiple comparison tests, the compact letter display (CLD) approach was used to categorize each group into statistically distinguishable subsets. Letters were assigned to indicate significance between groups. Groups with the same letter were not statistically different. Groups with non-overlapping letters were not statistically different. Analyses were performed using R version 4.4.1^60^. For comparison of only two groups, a Welch’s t-test with a significance level of α = 0.05 was used to identify groups with statistically different expression levels.

### Senescence

To check whether mutant plants have a senescence phenotype, individual 18-day-old plants regenerated from protoplasts were placed on the same plate containing 30 mL of PpNH4. Plates were left in the growth chamber for several weeks until at least one line turned brown (senescence). Senescence images were taken using a cellphone.

### Peptide synthesis and experiment

The PpPSY1 peptide (DY^S^EKPCANKKHDPSVCKNG) was obtained from Pacific Immunology (Ramona, CA, USA) and resuspended in double-distilled water. Peptide is tyrosine sulfated as indicated by (Y^S^).

To analyze bud and gametophore formation in response to peptide, moss lines were ground and immediately plated on cellophane-covered PpNH4 supplemented with 1µM PpPSY1. Tissue was moved to fresh peptide media weekly. After approximately 12 days, brightfield images were collected of each plate. At least 10 images were taken per plate. The experiment was repeated 3 independent times. The number of 3D structures per image was counted in each line. Data was normalized to the average number of 3D structures per image of the control condition for each line.

### RNA-seq analysis

Three-week-old ground tissue was flash frozen in liquid nitrogen and ground into powder using a mortar and pestle. Total RNA was extracted using RNAzol^61^. Specifically, powdered tissue was homogenized in RNAzol RT. Biphasic separation of RNA-rich supernatant and contaminating DNA, proteins, and polysaccharides was conducted with the addition of 4-bromoanisole. Total RNA was subsequently precipitated with isopropanol and Glycoblue and washed with 75% ethanol before recovered in nuclease-free water. Total RNA samples were treated with DNAse I using the Turbo DNA-*free* kit (Invitrogen) to remove contaminating genomic DNA. RNA quantity and quality was assessed by Azenta US, Inc. (Plainfield, NJ, USA) using the Agilent Tapestation System.

Library preparation, standard RNA-sequencing, and bioinformatics analysis was performed by Azenta US, Inc. Poly-A selection and External RNA Controls Consortium (ERCC) spike-in controls were added prior to library preparation. Standard RNA-sequencing was performed on an Illumina Novaseq platform using a 2×150 bp configuration to a depth of approximately 20 million paired-end (40 million total) reads per sample. Raw sequencing reads were cleaned using Trimmomatic v.0.36 then mapped to the Ppatens_318_v3.3_ERCC reference genome using the STAR aligner (v.2.5.2b). Gene-level hit counts were generated using featureCounts from the Subread package (v.1.5.2) and downstream differential expression analysis was performed using DESeq2.

Parameters for differentially expressed genes include False Discovery Rate (FDR, adjusted p-value) less than 0.05 and a log fold-change of 1, equal or greater than for up-regulated genes and equal or less than for down-regulated genes. Heatmaps were generated in RStudio (v2023.12.1+402) using the pheatmap R package^62^. Venn diagram was generated through InteractiVenn^63^. GO Term identification and analyses were done through Phytozome Biomart^59,64^. Identification of *P. patens* transcription factors was done through the Plant Transcription Factor Database^65^. Moss DEGs were mapped to *Arabidopsis thaliana* orthologs via Ensembl Plants BioMart: best-hit AGI, percent identity retained (plants.ensembl.org^66^). For the Heat map, Venn diagram, and GO Term analyses, only DEGs identified based on the above parameters were used. Published *Arabidopsis* DEG tables^24^ were used directly: *tri-1* (= *psyr1,2,3*) vs Col-0 root DEGs.

## Acknowledgements

We thank Sitwat Aman for advice and guidance with RNA isolation and RT-qPCR. We thank Valley Stewart for helpful discussions. This work was funded by the following grants: National Science Foundation (IOS-2436798) to MB and AMS, National Science Foundation (IOS-1954929) to PCR, National Institute of Health (R35 GM148173) to PCR, USDA-NIFA-AFRI Postdoctoral Fellowship (2023-67012-39889) to AMS, and Dartmouth College, Department of Biological Sciences (Class of 1978 Award) to DVT.

## Author Contributions

The study was conceived by DVT, AMS, SW, PCR, and MB. DVT and AMS designed, performed, and analyzed the experimental results. DVT, SW and MB designed the RNA-Seq study; AMS and ATAJ did the downstream RNA-Seq data analysis and visualization. DL performed and analyzed experimental results. The manuscript was written and edited by DVT, AMS, PCR, and MB with input from all authors.

## Data Availability

All data are available in the main text or in Supplementary Materials. The raw data for the RNA-Seq data has been submitted to the NCBI Gene Expression Omnibus (GEO).

