## Supplementary figures and images for "The Sulfated PSY Peptide Negatively Regulates Receptor Kinase Activity to Promote Growth"

### Fig. S1

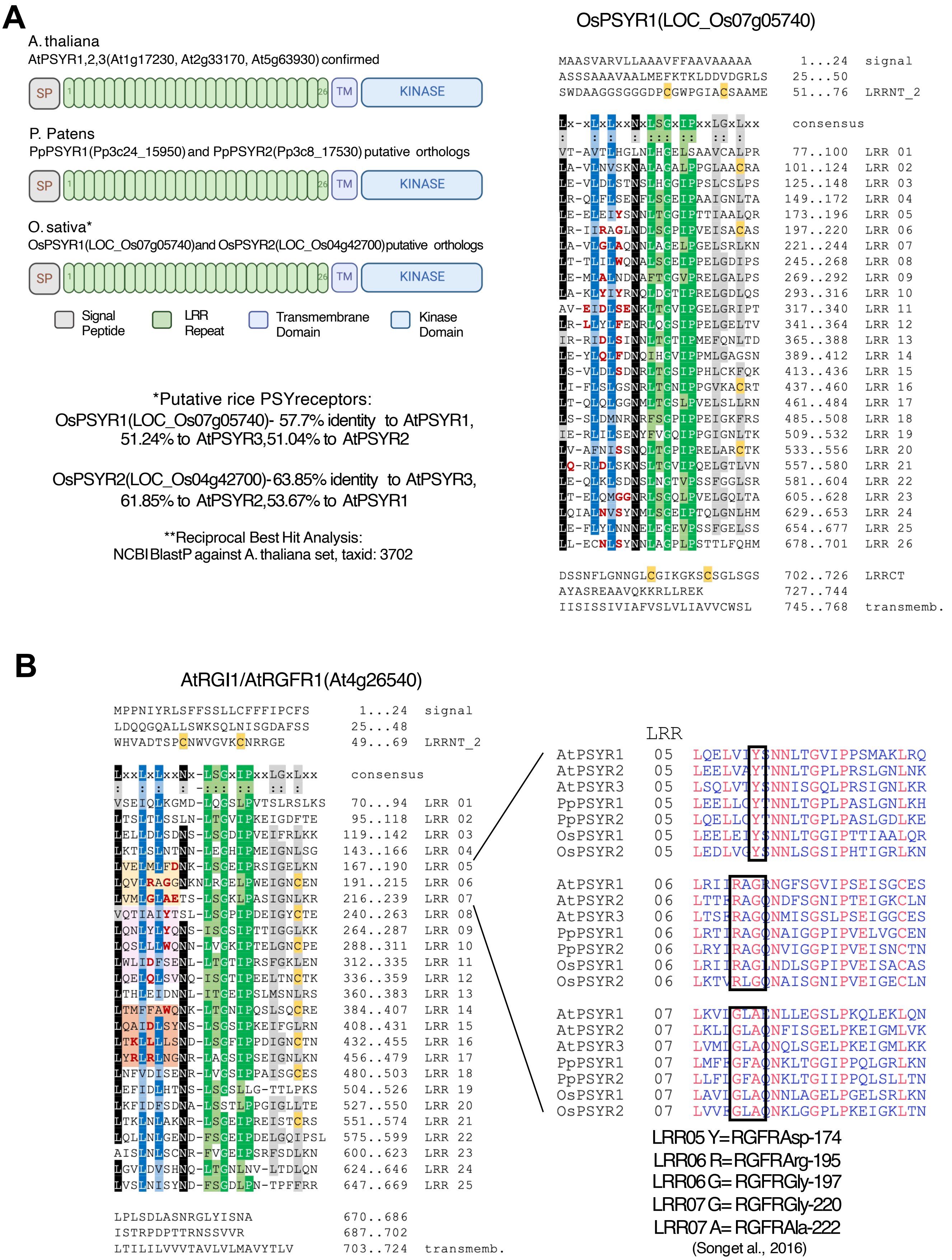

### Fig. S3

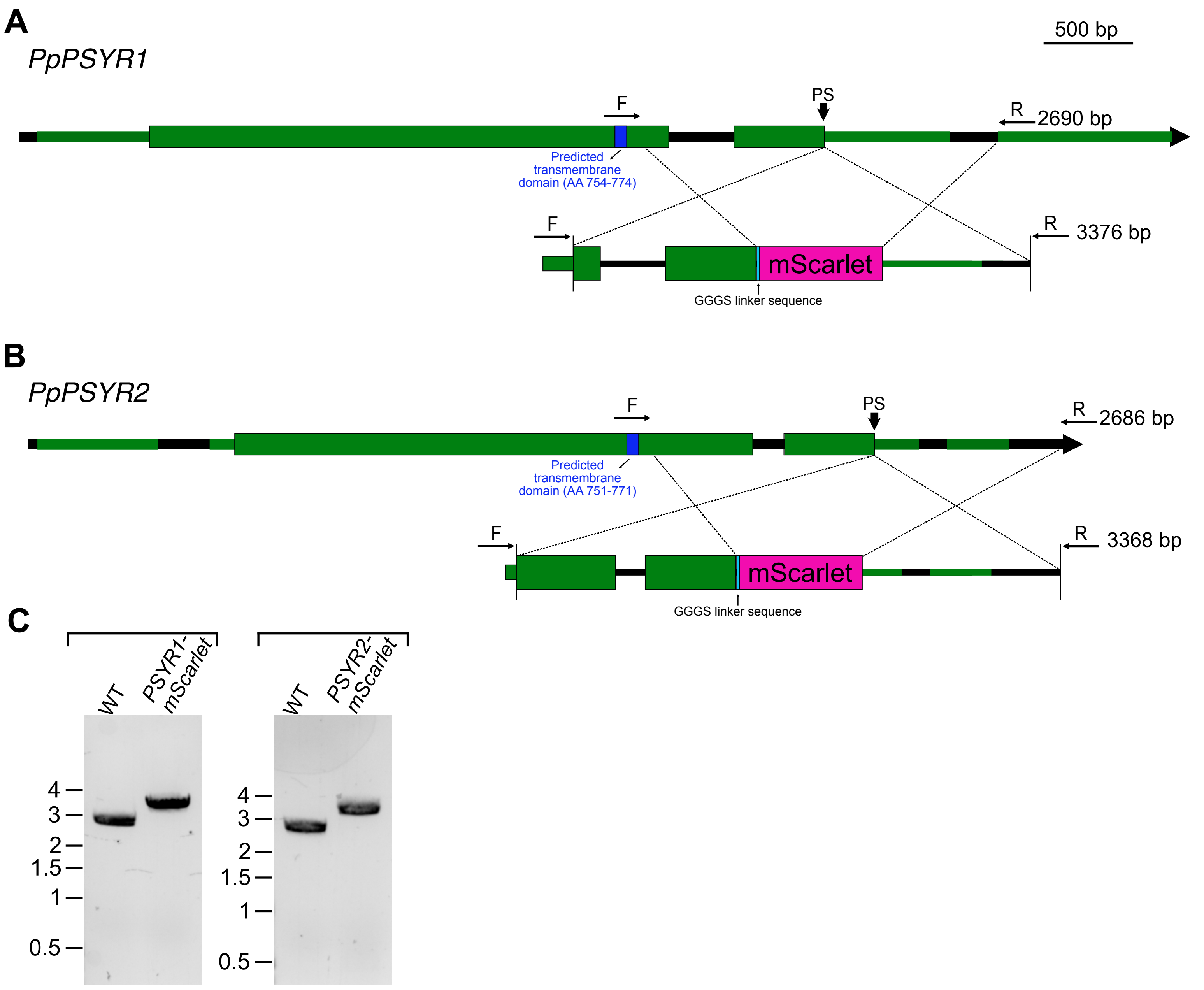

### Fig. S4

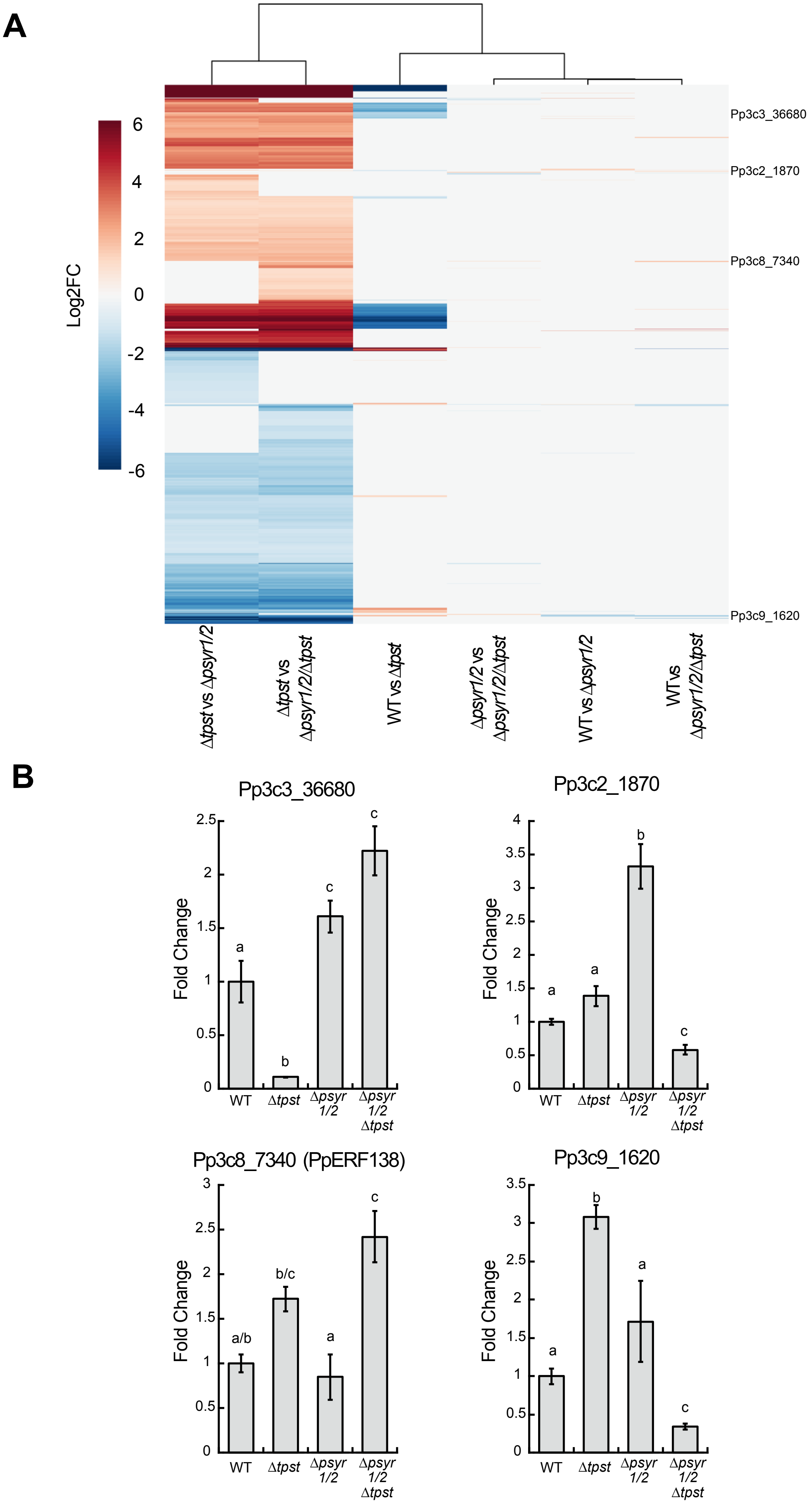

### Fig. S5

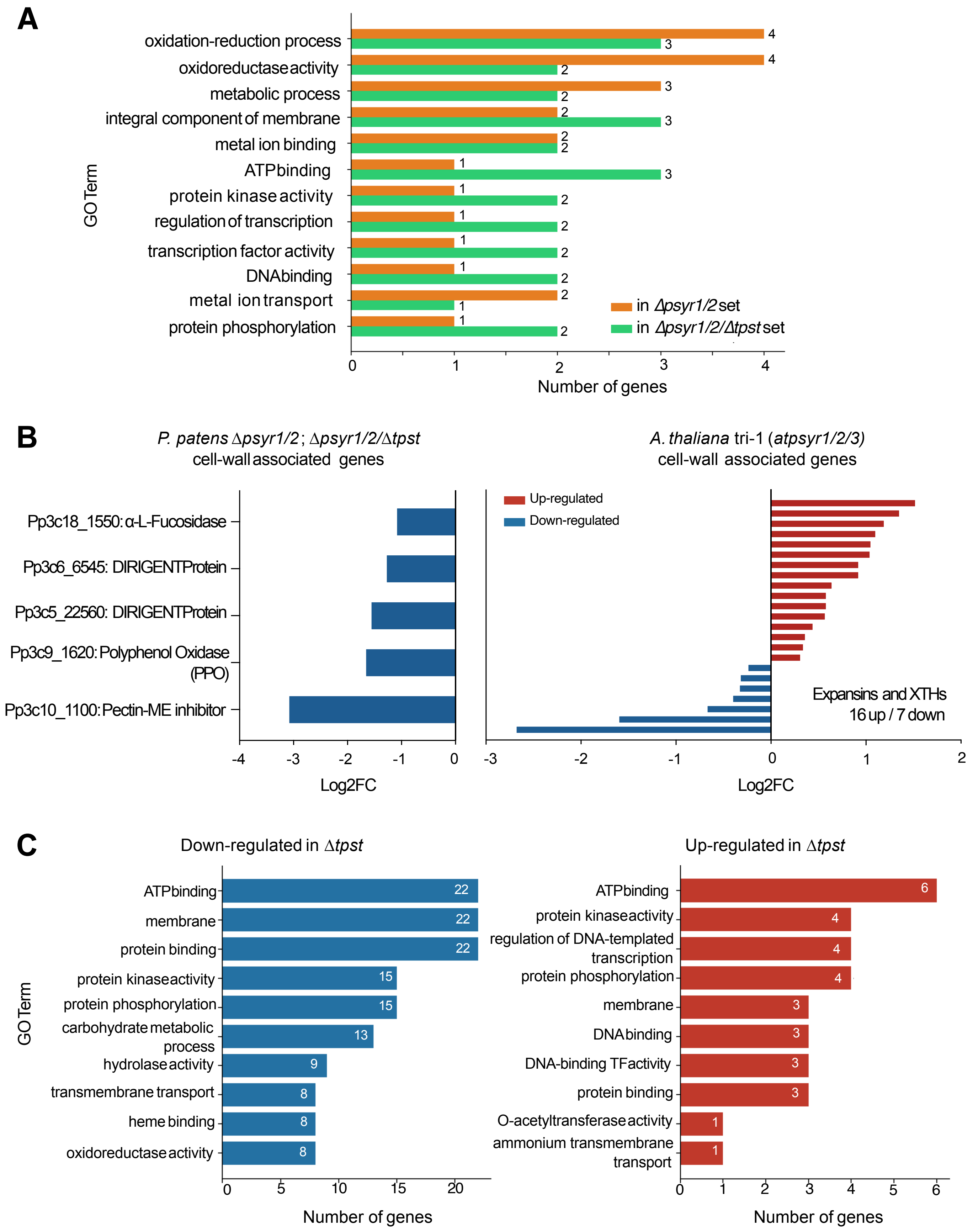

### Fig. S6

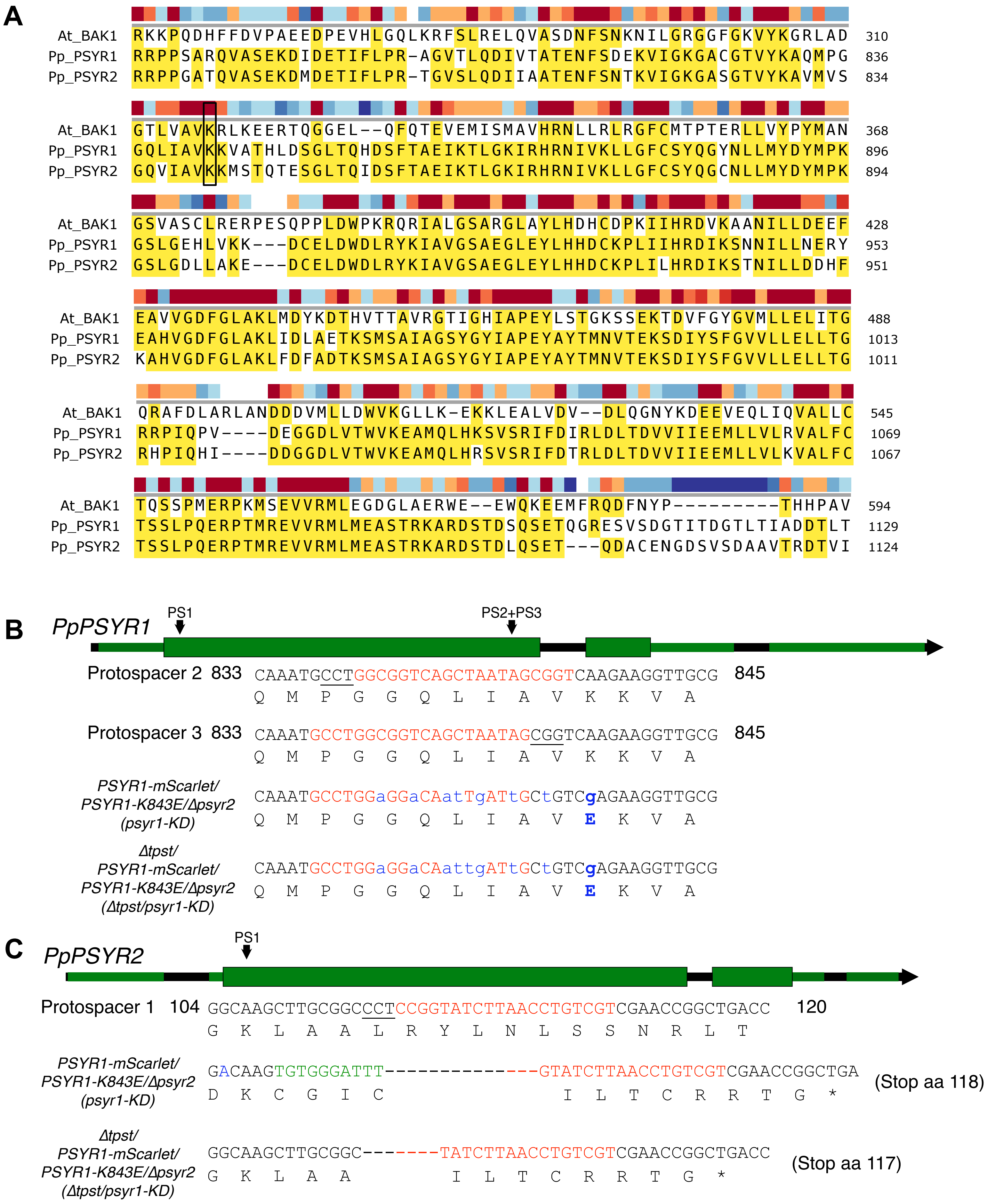

### Fig. S7

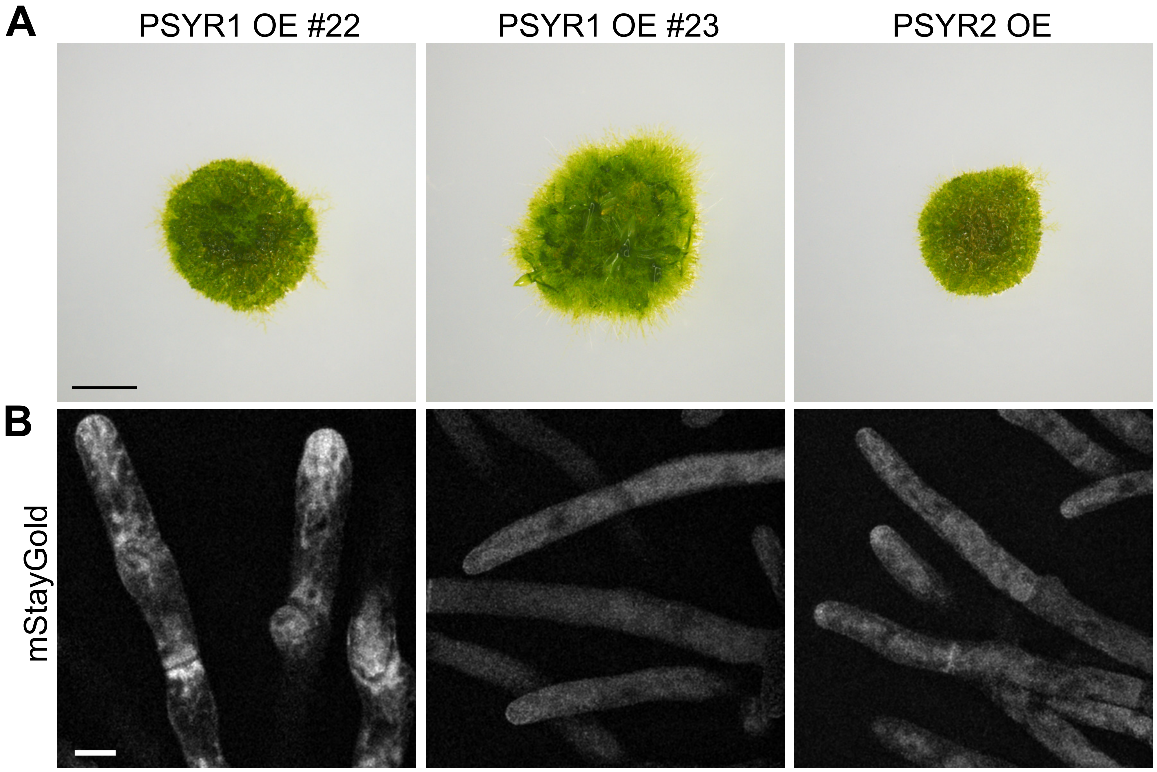


**Figure S7**

### Fig. S8

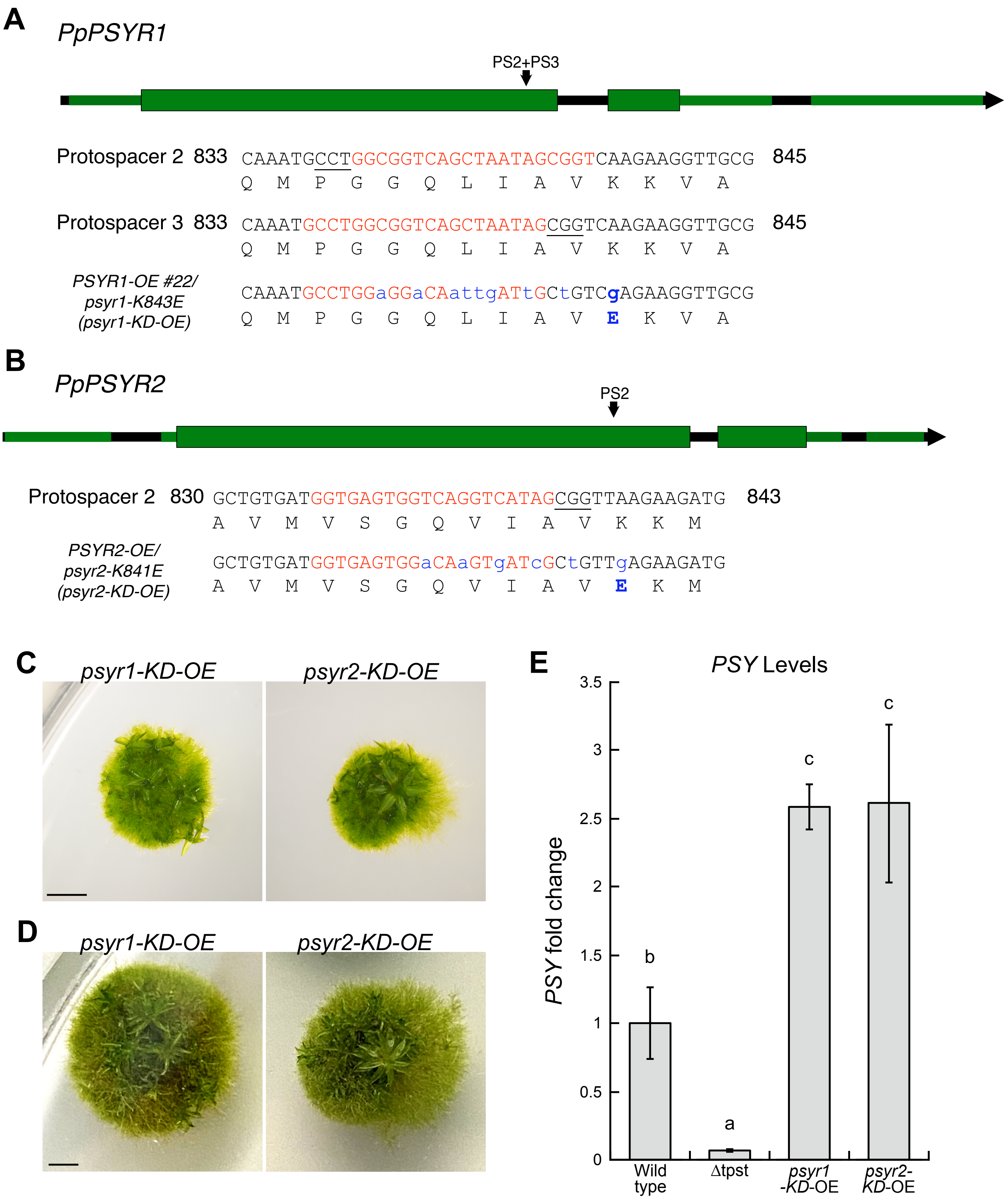
