## Supplementary material for "The Sulfated PSY Peptide Negatively Regulates Receptor Kinase Activity to Promote Growth": Fig. S2

**A** *PpPSYR1*

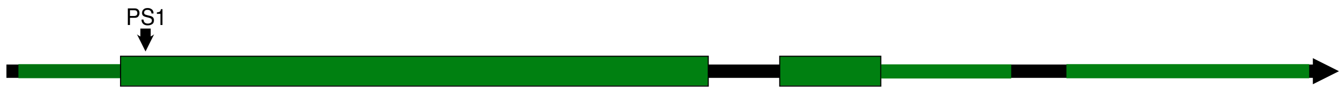


Protospacer 1 19 TTGCTCGGGTGGGCTCTCGCCGTGAGCTTAGTGGCGTTGCTG 32
L L G W A L A V S L V A L L

*Δpsyr2/Δpsyr1-14* *(Δpsyr1/2)* TTGCTCGGGTGGGCTCTCGCCGTG----------GCGTTGCTG (Stop aa 41)
L L G W A L A V A L L

33 TCGTGTCGGAGTTGCTGTGGGCTGTCCCCCAACTGA 41
S C R S C C G L S P N *

**B** *PpPSYR2*

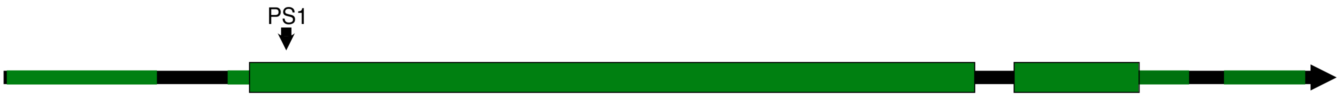


Protospacer 1 104 GGCAAGCTTGCGGCCCTCCGGTATCTTAACCTGTCGTCGAAC 117
G K L A A L R Y L N L S S N

*Δpsyr1/2* GGTTAAGCTT-------------------AACCTGTCGTCGTA (Stop aa 105)
G *

**C**

*PpTPST*

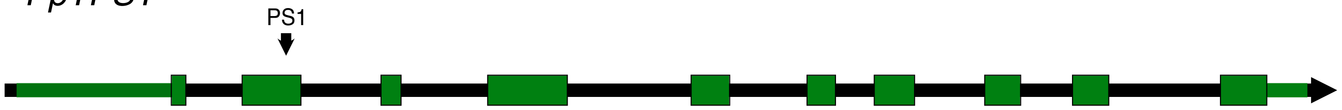


Protospacer 1 74 CTGTTCTTTTTGCATATTCCTCGAACCGGCGGCCGGACTTAT 87
L F F L H I P R T G G R T Y

*Δpsyr1/2/Δtpst-3,* *(Δpsyr1/2/Δtpst)* CTGTTCTTTTTGCATATTCCTCGAATAGCTGATTAAACTTAT (Stop aa 85)
L F F L H I P R I A D *

**Figure S2**
