## Supplementary Figure Legends for "The Sulfated PSY Peptide Negatively Regulates Receptor Kinase Activity to Promote Growth"

**Supplementary Figure 1. Conservation of LRR-RLK domains and predicted RGFR/RGI sulfotysorine residues.** A) LRR-RLK domain conservation between Arabidopsis PSY receptors with putative *P. patens* and *O. sativa* PSY Receptors (top left panel, LRR 1 and LRR26 numbered in LRR-domain). Figure made with BioRender.com. Reciprocal best hit analysis comparing putative OsPSYRs with AtPSYRs, indicating naming of OsPSYR1 and OsPSYR2 (bottom left panel). Sequence alignment of putative OsPSYR1(LOC_Os07g05740) ectodomain (Signal peptide, LRR-cap, LRR-domain to transmembrane domain). Residues highlighted in red are conserved in AtPSYRs, PpPSYRs, and OsPSYR2 (LOC_Os04g42700). B) Sequence alignment of the ectodomain (Signal peptide, LRR-cap, LRR-domain to transmembrane domain) for AtRGFR1/RGI3. Residues important for binding the RGF1 peptide are highlighted by the orange, purple, and red boxes. The residues important for recognition and binding of the sulfotyrosine residue of RGF1 is in the orange box (LRR05 to LRR07). The sequence alignments for AtPSYRs, PpPSYRs, and OsPSYRs for LRR05 and LRR07 are highlighted with the residues important for binding the sulfotyrosine listed (based on Song et la, 2016). Conserved residues with the AtPSYRs, PpPSYRs, and OsPSYRs are highlighted in red.

**Supplementary Figure 2. CRISPR-Cas9-mediated editing of *TPST, PSYR1,* and *PSYR2* loci.** Diagram of the *PSYR1* (A), *PSYR2* (B), and *TPST* (C) loci. Black lines denote intronic regions. Coding exons are represented by thick boxes. Green lines represent the 5' and 3' UTR. Location of the protospacers used to create the *psyr1*, *psyr2*, and *tpst* null mutants are denoted by PS and the corresponding sequence is represented by red text. The protospacer adjacent motif (PAM) is underlined. Dashed lines represent base pair deletions. Green text denotes base pair insertion. Blue text indicates a base pair change. All mutants have the indicated mutations resulting in early translational stops. A *tpst* null mutant was generating using the *psyr1* and *psyr2* null mutant described in (A) and (B) to create *Δpsyr1/2/Δtpst*.

**Supplementary Figure 3. Genotyping mScarlet tagging in the *PSYR* loci.** Diagram of the insertion of *mScarlet* into the locus of *PSYR1* (A) and *PSYR2* (B) via homology-directed repair. The locations of the protospacers (PS) are denoted by arrows. Thin black lines represent intronic regions. Thick green boxes denote coding exons. Thin green boxes represent the 5' and 3' UTR. The dashed vertical lines represent the junction between the knock-in construct and the genomic sequences located upstream and downstream of the insertion site. Arrows on the left and right side indicate forward and reverse primers for genotyping. The number adjacent to the reverse primer indicates the predicted PCR product size. C) PCR products obtained using the indicated primers in A and B. Product size denoted in kb.

**Supplementary Figure 4: Visualization of all differentially expressed genes across each experimental condition and RT-qPCR validation of candidate genes.** A) Hierarchically clustered heatmap of $\log_2$ fold-changes for the 2,237 genes differentially expressed in at least one comparison across the six comparisons; cell color indicates the $\log_2$ fold-change of the second-named genotype relative to the first (red = higher; blue = lower); white indicates no change or a gene not called differentially expressed in that comparison (value set to 0). Wild type (WT); *Δpsyr1/2* and *Δtpst/Δpsyr1/2* cluster together; Δ*tpst* is the outlier. B) RT-qPCR validation of RNA-Seq target genes highlighted in (B) in approximately 3- to 4-week-old ground tissue. Error bars indicate standard error (n=3). Different letters indicate significant differences determined by a one-way ANOVA with Tukey's multiple comparisons (α=0.05). Lists of all differentially expressed genes in (A) listed in Table S2.

**Supplementary Figure 5. Transcriptional responses across the receptor-null and sulfation-null comparisons.** A) Gene Ontology (GO) categories represented in both the *Δpsyr1/2* and *Δpsyr1/2/Δtpst* DEG sets, showing the number of genes contributed by each, and distinct genes converging on shared functions. B) Both PpPSY and AtPSY receptor-null systems converge on cell-wall loosening through opposite molecular strategies. Left, $log_2$ fold-change of wall-stiffening and phenolic-coupling enzymes in the moss receptor-null contrasts (*Δpsyr1/2* or (*Δpsyr1/2/Δtpst*). Right, $log_2$ fold-change of wall-loosening enzymes (expansins and xyloglucan endotransglucosylases/hydrolases) in *Arabidopsis tri-1 (atpsyr1,2,3/rek1,2,3)*, which are predominantly induced (16 up, red; 7 down, blue). Positive values denote up-regulation in the receptor-null genotype. *Arabidopsis* data from (*20*). C) Most frequent GO terms among genes down-regulated (B) and up-regulated (C) in *Δtpst* relative wild type (WT) (gene counts; descriptive, not tested against a genome-wide background). Lists of genes for each GO Annotation (A, B, C) highlighted in Table S2.

**Supplementary Figure 6. CRISPR-Cas9-mediated editing to generate the kinase-dead mutant lines.** A) Alignment of the kinase domains of Arabidopsis BAK1 (At_BAK1, aa251-594), PSYR1 (aa778-1129) and PSYR2 (aa776-1124). Black box highlights the position of the conserved Lysine317 in At_BAK1 used to determine the equivalent amino acid the *P. patens* PSY receptors. Diagram of the *PSYR1* (A) and *PSYR2* (B) loci in the *psyr1-KD* and *Δtpst/psyr1-KD* lines. Black lines denote intronic regions. Coding exons are represented by thick boxes. Green lines represent the 5' and 3' UTR. Location of the protospacer used to create *psyr1-K843E* via HDR and a *psyr2* null mutant by NHEJ is denoted by PS and the corresponding sequence is represented by red text. The PAM is underlined. Dashed lines represent base pair deletions. Green text denotes base pair insertion. Blue text indicates a base pair change. Lowercase text indicates a silent mutation. Blue capital letter represents the K843E point mutation in *PSYR1*.

**Supplementary Figure 7. *PSYR1* and *PSYR2* overexpression lines.** A) Brightfield images of 31-day-old plants regenerated from protoplasts. Scale bar, 0.2 cm. B) Maximum intensity projection images from confocal z-stacks of PSYR1-mStayGold or PSYR2-mStayGold signal in protonemata. Scale, 20 µm.

**Supplementary Figure 8. CRISPR-Cas9-mediated editing to generate *psyr1*-K843E and *psyr2*-K841E mutations.** Diagrams of the *PSYR1* (A) and *PSYR2* (B) loci in the *psyr-KD-OE* lines. Black lines denote intronic regions. Coding exons are represented by thick boxes. Green lines represent the 5' and 3' UTR. Location of the protospacer used to create the point mutations via HDR is denoted by PS and the corresponding sequence is represented by red text. The PAM is underlined. Dashed lines represent base pair deletions. Blue text indicates a base pair change. Lowercase text indicates a silent mutation. Blue capital letter represents a point mutation. C) Bright field images of 31-day-old plants regenerated from protoplasts. Scale, 0.2 cm. D) The same plants depicted in C at 50-days-old. Scale, 0.2 cm. D) *PSY* expression levels in approximately 3-week-old tissue. Significant differences determined by a one-way ANOVA with Tukey's multiple comparisons (α=0.05) are indicated by different letters.
