## Supplementary material for "The Sulfated PSY Peptide Negatively Regulates Receptor Kinase Activity to Promote Growth": Table S1

**Table S1**. List of mutant lines used and created.

| Name | Background | Genetic alteration | Translation |
|---|---|---|---|
| Wild type | Gransden 2004 (*67*) | N/A | N/A |
| *Δtpst* (*26*) | Wild type | 7 bp deletion | Stop at AA90, aa 81-89 are irrelevant |
| *Δpsyr1/2* | Wild type | *PSYR1:* 9 bp deletion, 4 bp insertion | *PSYR1:* Stop at AA41, aa 40 is irrelevant, deletion of AA 27-29 |
| | | *PSYR2:* 1 bp insertion, 1 bp change, 18 bp deletion | *PSYR2*: Stop at AA105, silent mutation in aa 104 |
| *Δpsyr1/2/Δtpst* | *Δpsyr1/2* | Stop oligo insertion | Stop at AA85, aa 82-84 are irrelevant |
| *PSYR1-mScarlet* | Wild type | Insertion of the sequence encoding for mScarlet at the 3' end of the *PSYR1* gene at the endogenous locus | C-terminal fusion of PSYR1 with mScarlet |
| *Δtpst/PSYR1-mScarlet* | *Δtpst* | Insertion of the sequence encoding for mScarlet at the 3' end of the *PSYR1* gene at the endogenous locus | C-terminal fusion of PSYR1 with mScarlet |
| *PSYR2-mScarlet* | Wild type | Insertion of the sequence encoding for mScarlet at the 3' end of the *PSYR2* gene at the endogenous locus | C-terminal fusion of PSYR2 with mScarlet |
| *Δtpst/PSYR2-mScarlet* | *Δtpst* | Insertion of the sequence encoding for mScarlet at the 3' end of the *PSYR2* gene at the endogenous locus | C-terminal fusion of PSYR2 with mScarlet |
| *psyr1-K843E/Δpsyr2* (*psyr1-KD*) | *PSYR1-mScarlet* | *PSYR1:* A5109G (AA change), C5090A(silent), T5093A (silent), G5096A (silent), C5097T (silent), A5099G (silent), A5102T (silent), G5105T (silent)<br>*PSYR2:* 1 bp change, 10 bp deletion | *PSYR1:* K841E (kinase inactive)<br>*PSYR2*: Stop at AA118, aa 104-117 are irrelevant |
| *Δtpst/psyr1-K843E/Δpsyr2* (*Δtpst/psyr1-KD*) | *Δtpst/PSYR1-mScarlet* | *PSYR1:* A5109G (AA change), C5090A(silent), T5093A (silent), G5096A (silent), C5097T (silent), A5099G (silent), A5102T (silent), G5105T (silent)<br>*PSYR2*: 7 bp deletion | *PSYR1:* K841E (kinase inactive)<br>*PSYR2*: Stop at AA117, aa 109-116 are irrelevant |

| | | | |
|---|---|---|---|
| *ACT1::PSYR1-mStayGold* #22 and #23 (*PSYR1-OE #22 and #23*) | *PSYR1-mScarlet* | Insertion of the sequence encoding for mStayGold at the 3' end of the *PSYR1* CDS targeted to the pTK locus | Overexpression of C-terminal fusion of PSYR1 with mStayGold |
| *ACT1::PSYR2-mStayGold #9* (*PSYR2-OE*) | *PSYR1-mScarlet* | Insertion of the sequence encoding for mStayGold at the 3' end of the *PSYR2* CDS targeted to the pTK locus | Overexpression of C-terminal fusion of PSYR2 with mStayGold |
| *PSYR1-OE #22/psyr1-K843E* | *PSYR1-OE #22* | A5109G (AA change), C5090A(silent), T5093A (silent), G5096A (silent), C5097T (silent), A5099G (silent), A5102T (silent), G5105T (silent) | K843E (kinase inactive) |
| *PSYR2-OE /psyr2-K841E* | *PSYR2-OE* | A5518G (AA change), T5502A (silent), G5505A (silent), C5508G (silent), A5511C (silent), G5514T (silent), | K841E (kinase inactive) |
