## Supplementary material for "The Sulfated PSY Peptide Negatively Regulates Receptor Kinase Activity to Promote Growth": Table S3

**Table S3.** Table of oligos

| Name | Sequence (5' to 3') | Purpose |
|---|---|---|
| PSYR1-PS1 | ATAGGAAGGTGGAGGACTTACAACCCATGGCTCTCGCCGTGAGCTTAGgttttagagctatgctgaaaag | PSYR1 protospacer |
| PSYR1-gt-F | GGAGTGGTTTGTCGACGAAGG | PSYR1 genotyping |
| PSYR1-gt-R | CTGACCAGCTCGAATTGTCCG | PSYR1 genotyping |
| PSYR1-gt-R3 | GACAGCAACGCCACTAAGC | PSYR1 genotyping, anneals to protospacer 1 |
| PSYR2 PS2 | ATAGGAAGGTGGAGGACTTACAACCCATACGACAGGTTAAGATACCGGgttttagagctatgctgaaaag | PSYR2 protospacer |
| PSYR2-gt-F1 | CATAGCTCGGCATGAGGAGG | PSYR2 genotyping |
| PSYR2-gt-R2 | GGAGGTTAGGTTGCCGAGAC | PSYR2 genotyping |
| PSYR2-gt-R1 | ACGACAGGTTAAGATACCGGAGG | PSYR2 genotyping, anneals to protospacer 1 |
| TPST protospacer 1 | ATAGGAAGGTGGAGGACTTACAACCCATatattcctcgaaccggcggcGTTTTAGAGCTATGCTGAAAAG | TPST protospacer |
| TPST stop oligo F | ggaaaattctggaacttctgttctttttgcatattcctcgaaTAGCTGATTAAacttatcaccagtggtatggatttact | Stop oligo for TPST |
| TPST stop oligo R | agtaaatccataccactggtgataagtTTAATCAGCTAttcgaggaatatgcaaaaagaacagaagttccagaattttcc | Stop oligo for TPST |
| TPST-comp-F | ttctctcttgtctctgttgatcacc | TPST genotyping |
| TPST-cutsiteF | atattcctcgaaccggcgg | TPST genotyping, anneals to protospacer 1 |
| TPST-compR | gcgcaacagtctacgtgc | TPST genotyping |

| | | |
|---|---|---|
| PSYR1_5arm_F | ACAAGTTTGTACAAAAAAGCAGGCTCATTCATCGCGACATCAAGTCCAAC | PSYR1-mScarlet construct preparation |
| PSYR1_5arm_R | tgctcaccatcgatcctccgccaccTGGTTTATCAGAGTTGGTCTTCCCAG | PSYR1-mScarlet construct preparation |
| PSYR1_3arm_F | cggcatggacgagctgtacaagTGAAGGGCAAGGCAGGAATGG | PSYR1-mScarlet construct preparation |
| PSYR1_3arm_R | ACCACTTTGTACAAGAAAGCTGGGCCATCAACATCCATCCATACCTGAAA | PSYR1-mScarlet construct preparation |
| PSYR2_5arm_F | ACAAGTTTGTACAAAAAAGCAGGCTGTATGATTACATGCCGAAGGGGA | PSYR2-mScarlet construct preparation |
| PSYR2_5arm_R | cttgctcaccatcgatcctccgccaccAGGCTTGTCAGAATTCTCAGAAATGATC | PSYR2-mScarlet construct preparation |
| PSYR2_3arm_F | cggcatggacgagctgtacaagTAGCTCGGCAGGAAAGGGCTACT | PSYR2-mScarlet construct preparation |
| PSYR2_3arm_R | ACCACTTTGTACAAGAAAGCTGGGGCGCACAACTACTCGAGTACC | PSYR2-mScarlet construct preparation |
| GGGS-FP-F | ggtggcggaggatcgatggtgagcaagggcgagg | GGGS linker sequence for mScarlet |
| FP-no stop-R | cttgtacagctcgtccatgcc | mScarlet amplification |
| PSYR1 PS2 | ATAGGAAGGTGGAGGACTTACAACCCATGCCTTGCCCTTGTACCAGGGgttttagagctatgctgaaaag | PSYR1 c-terminus tagging protospacer |
| PSYR2 PS2 | ATAGGAAGGTGGAGGACTTACAACCCATCCTGCCGAGTGATTGTACCAgttttagagctatgctgaaaag | PSYR2 c-terminus tagging protospacer |
| PSYR1-gt-F2 | CCGCCACCGAAAATTTCAGTGAT | PSYR1-genotyping |
| PSYR1-gt-R3 | ACCGCTGTGTGAGTACGTAACA | PSYR1-genotyping |
| PSYR2-gt-F2 | CTGGCGCTACACAAGTAGCAAG | PSYR2-genotyping |
| PSYR2-gt-R3 | TGTTGCTGTAAACGTATACCCGGA | PSYR2-genotyping |
| PSYR1-ps3 | ATAGGAAGGTGGAGGACTTACAACCCATACCGCTATTAGCTGACCGCCgttttagagctatgctgaaaag | PSYR1 protospacer for PSYR1-K843E |

| | | |
|---|---|---|
| PSYR1-ps4 | ATAGGAAGGTGGAGGACTTACAACCCATGCCTGGCGGTCAGCTAATAGgttttagagctatgctgaaaag | PSYR1 protospacer for PSYR1-K843E |
| PSYR1_K843E_oligo-F | GCATGTGGGACAGTGTACAAAGCGCAAATGCCTGGaGGaCAattgATtGCtGTCgAGAAGGTTGCGACTCATCTAGACTCAGGTTTGACT | PSYR1 kinase dead oligo |
| PSYR1_K843E_oligo-R | AGTCAAACCTGAGTCTAGATGAGTCGCAACCTTCTcGACaGCaATcaatTGtCCtCCAGGCATTTGCGCTTTGTACACTGTCCCACATGC | PSYR1 kinase dead oligo |
| PSYR1-K843E-gt-F1 | CCAGTTCCAGTAGCATGTCCAC | PSYR1 genotyping |
| PSYR1-K843E-gt-R1 | TGACCGCTATTAGCTGACCG | PSYR1 genotyping, anneals to protospacer |
| PSYR1-K843E-gt-R2 | ACCCTGCTATTGCAGACATCG | PSYR1 genotyping |
| PSYR2-ps2 | ATAGGAAGGTGGAGGACTTACAACCCATGGTGAGTGGTCAGGTCATAGgttttagagctatgctgaaaag | PSYR2 protospacer for PSYR2-K841E |
| PSYR2_K841E_oligo-F | AGTGGAACAGTCTACAAAGCTGTGATGGTGAGtGGaCAaGTgATcGCtGTTgAGAAGATGTCCACTCAGACAGAATCAGGCTTGACTCAG | PSYR2 kinase dead oligo |
| PSYR2-K841E_oligo-R | CTGAGTCAAGCCTGATTCTGTCTGAGTGGACATCTTCTcAACaGCgATcACtTGtCCaCTCACCATCACAGCTTTGTAGACTGTTCCACT | PSYR2 kinase dead oligo |
| PSYR2-K841E-gt-F | GTGGTCAGGTCATAGCGGTTAAG | PSYR2 genotyping, anneals to protospacer |
| PSYR2-K841E-gt-R | CACAGCGATCTTATATCGCAGGTC | PSYR2 genotyping |
| PSYR2-F2 | CTGGCGCTACACAAGTAGCAAG | PSYR2 genotyping |
| attB1-F | ACAAGTTTgtacaaaaaagcaggct | Linearize pTK-UBI plasmid around Ubi fragment |
| pTK-5arm-R | TCGACTTCCCTGCTTCGATCC | Linearize pTK-UBI plasmid around Ubi fragment |
| Act1pro-F | GGATCGAAGCAGGGAAGTCGAtcgaggtcattcatatgcttgagaagag | Act1 promoter amplification for pTK-Act1 |
| Act1pro-R | agcctgcttttttgtacAAACTTGTvggatcctctagatcttctacctacaaaaaagc | Act1 promoter amplification for pTK-Act1 |

| | | |
|---|---|---|
| PSYR1-cDNA-F | ATGACGAGGGGGAGGGAGAC | PSYR1 cDNA amplication |
| PSYR1-cDNA-R | TGGTTTATCAGAGTTGGTCTTCCCAG | PSYR1 cDNA amplication |
| attB1-PSYR1-F | ACAAGTTTgtacaaaaaagcaggctATGACGAGGGGGAGGGAGAC | PSYR1 fragment for pTK-Act1 |
| PSYR1-linker-R | tcctgaggctcccgatgctccTGGTTTATCAGAGTTGGTCTTCCCAG | PSYR1 fragment for pTK-Act1 |
| linker-mStayGold | ggagcatcgggagcctcaggagcatcgatggtgtctacaggcgaggag | mStayGold fragment for pTK-Act1 |
| mStayGold-attB2-R | ACCACTTTGTACAAGAAAGCTGGGTttacaggtgggcctccagggtctc | mStayGold fragment for pTK-Act1 |
| attB1-PSYR2-F | ACAAGTTTgtacaaaaaagcaggctATGAGGAGGAGAGGGGCGTGG | PSYR2 fragment for pTK-Act1 |
| PSYR2-linker-R | tcctgaggctcccgatgctccAGGCTTGTCAGAATTCTCAGAAATGATC | PSYR2 fragment for pTK-Act1 |
| mStayGold-R2 | ctggtgcacacgtacttgcc | PSYR1 or PSYR2-mSG genotyping |
| EF1alpha-RT-F2 | CCCTTCAGGATGTCTACAAGATT | RT-RT-qPCR primer |
| EF1alpha-RT-R2 | CCCTTCAGGATGTCTACAAGATT | RT-qPCR primer |
| PSYR1-RT-F | GGTGGTGCTTTGTTGATGATATT | RT-qPCR primer |
| PSYR1-RT-R | TCGTCTCGTCTATGTCCTTCT | RT-qPCR primer |
| PSYR2-RT-F | GCCTCTTTGCGAAGCTAAATG | RT-qPCR primer |
| PSYR2-RT-R | AGCCATCGGTGTTGGTAAG | RT-qPCR primer |
| PSY-RT-F | TTCCAGCAAGAGAAGGTTGG | RT-qPCR primer |
| PSY-RT-R | CTCCACGAGATTTGCCTCAG | RT-qPCR primer |
| Pp3c2_1870-RT-F | GCGCGAGTGGTAGTAATGAA | RT-qPCR primer |
| Pp3c2_1870-RT-R | ACTGATCTGCGGGCTTTAAT | RT-qPCR primer |
| Pp3c3_36680-RT-F | GATGTTAAGGCTGCCAACATTC | RT-qPCR primer |
| Pp3c3_36680-RT-R | GTGACATGGGTGTCCTTATAGTC | RT-qPCR primer |
| PpERF138-RT-F | AATGGCAGCTAGAGCGTATG | RT-qPCR primer |

| | | |
|---|---|---|
| PpERF138-RT-R | AGTGGCATTCAGGAAGTGATT | RT-qPCR primer |
| Pp3c9_1620-RT-F | CACGAATACCGACCGACTTT | RT-qPCR primer |
| Pp3c9_1620-RT-R | CCTCGTCATAGAACGTGAACTC | RT-qPCR primer |
